# Patterns of phenotypic plasticity among populations of three Mediterranean pine species and implications for evolutionary responses to climate change

**DOI:** 10.1101/716084

**Authors:** Natalia Vizcaíno-Palomar, Bruno Fady, Ricardo Alía, Annie Raffin, Sven Mutke, Marta Benito Garzón

## Abstract

**Aim:** Under rapid environmental change, phenotypic plasticity, if adaptive, could increase the odds for organisms to persist. Environmental variation over time is an important source of phenotypic plasticity. Likewise, phenotypic plasticity can vary with age in many organisms. However, little is known on phenotypic plasticity variation across species’ ranges. Our aims are: (i) to assess whether populations’ phenotypic plasticity is related to the inter-annual climate variation under which populations have evolved during the last century; (ii) to compare phenotypic plasticity among developmental classes; and (iii) to predict phenotypic plasticity across’ species ranges.

**Location:** Europe and North-Africa.

**Time period:** 1901-2014.

**Major taxa studied:** *Pinus nigra, P. pinaster* and *P. pinea*.

**Methods:** We used 372 646 individual tree height measurements at three developmental classes from a wide network of 38 common gardens in Europe and North Africa with provenances covering the distribution range of the species. With this data, we: i) build linear mixed-effect models of tree height as a function of tree age, population and climate; ii) estimate populations’ reaction norms from the fitted models; iii) calculate populations’ phenotypic plasticity indexes; iv) build models of populations’ phenotypic plasticity indexes as a function of inter-annual climate variation during the last century.

**Results:** We found that i) most populations that have evolved under high inter-annual climate variation, in either maximum or minimum values in temperature or precipitation, exhibited high values of plasticity in tree height; ii) phenotypic plasticity for tree height was higher in young trees than in older ones, iii) phenotypic plasticity did not follow any particular geographical pattern across species’ ranges.

**Main conclusions:** Phenotypic plasticity across the three Mediterranean pines’ ranges is related with the climate variation experienced over time and calls into question whether this plasticity could be adaptive and hence beneficial to cope with climate change in the short-term.

## Introduction

Climate change is reshuffling species distribution ranges from marine to terrestrial systems, altering current ecosystems functioning and structure through disruption of species interactions at temporal and/or spatial scales (Lenoir *et al*., 2008; Poloczanska *et al*., 2013). To survive under new climates, organisms can move to more favorable environments (Chen *et al*., 2011; Sunday *et al*., 2011), or persist *in-situ* by changes in their genetic composition or adjusting to environmental changes using phenotypic plasticity (West-Eberhard, 2003; Pulido & Berthold, 2004). Evolutionary responses to climate change will imply changes in allele frequencies that need many generations to arise (Bradshaw & Holzapfel, 2001; Reale *et al*., 2003; Franks *et al*., 2007), whereas plastic responses can occur without changes in the genetic structure (Sultan, 2000; Valladares *et al*., 2014) within one generation (or even longer when including trans-generational effects, Donelson *et al*., (2018)). Thus, phenotypic plasticity can provide a rapid response, whereas evolutionary responses need longer time depending on the lifespan of organisms. For the particular case of trees, with very long generation times and large gene flow among populations, genetic adaptation occurs at long time scales (Savolainen *et al*., 2007). For example, evolutionary adjustments to match new climates could need more than 1500 years in *Pinus sylvestris* (Rehfeldt *et al*., 2002). Therefore, plasticity is often the main mechanism for tree populations to respond *in-situ* to rapid climate change (Benito Garzón *et al*., 2019).

Environmental variation, either spatial or temporal, may promote differentiation in phenotypic plasticity among populations (Vizcaíno-Palomar *et al*., 2016). In this context, some studies have shown that more plastic genotypes are promoted under greater heterogeneity (Lind & Johansson, 2007; Canale & Henry, 2010; Baythavong, 2011; Lázaro-Nogal *et al*., 2015). However, phenotypic plasticity may not be always advantageous, and sometimes it can be detrimental. For example, high values of plasticity can be associated with low values of fitness-related traits as survival, biomass, or reproduction (e.g. Sánchez-Gómez *et al*., (2006); Molina-Montenegro & Naya, (2012)). Likewise, changes in plasticity can occur during the lifespan of organisms due to morphological and physiological adjustments to the environment (Evans, 1972; Coleman *et al*., 1994; Mitchell & Bakker, 2014). Hence, we could expect to find differences in phenotypic plasticity for fitness-related traits between early and mature stages of development (Mediavilla & Escudero, 2004). For instance, we can expect high plasticity in seedlings that present small root systems located in the shallow soil layers with great variation in soil moisture in contrast to mature trees with deep root systems reaching to more stable layers of soil moisture over the year. Hence, greater plasticity at the recruitment stage could be favorable for plant establishment in the community. Taken altogether, we could expect that phenotypic plasticity can vary across the species’ distribution ranges and within-species lifespan.

The complex topography and orography of the Mediterranean basin, with its inter and intra-seasonal climate variation, and its recent story of species’ expansions from refugia after the Last Glacial Maximum could have promoted differentiation in phenotypic plasticity among populations across species’ ranges (Médail & Diadema, 2009). As a result, Mediterranean pine species present patchy distributions with differentiated patterns of genetic diversity and local adaptation, reviewed in Fady, (2012). For instance, although *P. nigra* has a larger distribution than *P. pinaster*, both present moderate-high population differentiation in neutral genetic diversity (Soto *et al*., 2010) and in quantitative traits, such as tree height, diameter, height-diameter allometry, survival, etc. (Varelides *et al*., 2001; Taïbi *et al*., 2016; Vizcaíno-Palomar *et al*., 2016). On the contrary, *P. pinea* presents very low levels in genetic diversity across its range (Vendramin *et al*., 2008), as well as low differentiation for quantitative traits, such as in tree height (Mutke *et al*., 2010, 2013; Sánchez-Gómez *et al*., 2011).

Assessing populations’ phenotypic plasticity responses across the species’ ranges requires the use of phenotypic data measured from multiple common gardens, ideally a minimum of three (Arnold *et al*., 2019), installed across large environmental gradients in which a suite of populations from varied origins are planted. These experimental designs permit to fit populations’ non-linear phenotypic responses to the environment, known as ‘reaction norm’ curves (Gavrilets & Scheiner, 1993; Schlichting & Pigliucci, 1998), from which quantifying populations’ phenotypic plasticity is straightforward (Arnold *et al*., 2019). Furthermore, populations’ phenotypic responses can be used to quantify populations’ phenotypic plasticity using indices (Valladares *et al*., 2006).

In this study, we used tree height, a fitness-related trait (King, 1990; Savolainen *et al*., 2007), measured in a wide network of common gardens established in Europe and North Africa for *Pinus nigra*, *P. pinaster* and *P. pinea* (Vizcaíno-Palomar *et al*., 2019). We fitted linear mixed-effect models of tree height to: i) assess whether populations’ phenotypic plasticity is related to the inter-annual climate variation under which populations have evolved during the last century; ii) compare phenotypic plasticity among developmental classes; iii) predict phenotypic plasticity across species ranges.

## Material and methods

### Provenance trials, species and phenotypic data

We used tree height recorded in common garden networks for three pine species: *Pinus nigra* Arn., *P. pinaster* Aiton and *P. pinea* L. (see Figure S1 in Supporting Information). For *Pinus nigra*, we used 192 221 measurements of individual tree height recorded in 15 trials distributed across three countries (France, Germany and Spain) where 78 populations (provenances) from origins covering the entire range of the species were planted. Trials were planted between years 1968 and 2009 and tree heights were measured between 2 and 18 year-old. For *P. pinaster* we used 123 801 measurements of individual tree height recorded in 14 trials established across three countries (France, Morocco and Spain) and 182 populations covering the entire range of the species. Trials were installed between years 1966 and 1992 and tree heights were measured between 2 and 34 year-old. For *P. pinea,* we used 56 624 measurements of individual tree height recorded in 9 trials established in France and Spain, where a total 55 populations covering the entire range of the species were planted. Trials were established between years 1993 and 1997, and tree heights were measured between 2 and 22 year-old. Further description of these databases can be found in Vizcaíno-Palomar *et al*., (2019).

To analyse the effect of age on phenotypic plasticity we defined three developmental classes (DC. 1, DC. 2 and DC. 3) covering the range of ages of each species. In all species, DC.1 included information for 4 year-old trees, DC.2 included trees of 8, 13, and 9 year-old, and DC.3 included information for 14, 24, and 22 year-old trees, for *P. nigra, P. pinaster* and *P. pinea*, respectively.

### Climate data

We used the EuMedClim dataset that provides annual measurements between 1901 and 2014, at 30 arc-seconds (∼ 1km) of resolution (http://gentree.data.inra.fr/climate/datasets/; Fréjaville & Benito Garzón, (2018)). We used a total of 21 climatic variables related with either annual or seasonal parameters of climate in terms of precipitation and temperature (Table S1). From this database, we computed the following climate-related variables and indices:

i. Long-term climate effect on trees’ height population (clim_p_) was calculated as the average climate at the population origin between the beginning of the 20^th^ century (1901) and the year before the trees were planted in the trials. This ‘long-term’ effect reflects the climate that occurred when the planted seeds were generated, and it can be related to population effects.
ii. Short-term climate effect on trees’ height population (clim_t_) was calculated as the average climate at the trial of the last 3 years including the year when measurements were taken. This definition assures that plastic responses are measured under equal periods of time in all trees for the three species, easing comparisons. This ‘short-term’ effect was defined to reflect the plastic response of tree height to recent climate.
iii. Inter-annual climate variation indices during the 20^th^ at the population origin. We computed the standard deviation (sd) of seven climate variables selected to reflect the past climate variation encountered by the tree populations since the beginning of 20^th^ century (1901) and the year before the trees were planted. Specifically, we computed the standard deviation (sd) of the mean annual temperature (sd bio1), sd of the mean diurnal temperature range (sd bio2), sd of the maximum temperature of the warmest month (sd bio5), sd of the minimum temperature of the coldest month (sd bio6), sd of the annual precipitation (sd bio12), sd of the precipitation of the wettest month (sd bio13) and sd of the precipitation of the driest month (sd bio14).

All climate-related variables and indices were standardized for further analyses.

### Statistical analyses

We used linear mixed-effect models to account for the following effects: tree age, genetics (approached by the climate at the population origin, clim_p_) and plasticity (approached by the climate at the trial, clim_t_) on tree height measured across the networks of provenance tests for the three species. Afterwards, we predict populations’ phenotypic responses across the climatic range covered by the trials to compute phenotypic plasticity indices to summarize these curves. Our approach allowed us to obtain linear or non-linear curve responses as these are very common in nature (Arnold *et al*., 2019). Then, phenotypic plasticity values can be estimated with more accurate.

#### 1. Linear mixed-effect models of tree height responses accounting for age and climate

For each species, we selected one climate variable for the population (hereafter clim_p_) and another for the trial (hereafter clim_t_). This selection was based upon the complementary use of linear mixed-effects models and on principal components analyses (PCA) of the climate variables (see Appendix S1 for a detailed description). For *P. nigra,* we selected mean annual temperature (bio1) for clim_t_ and annual water availability (WAI) for clim_p_. For *P. pinaster,* annual potential evapotranspiration (PET) for clim_t_ and winter precipitation (prec.djf) for clim_p_. And for *P. pinea,* maximum temperature of the warmest month (bio5) for clim_t_ and summer precipitation (prec.jja) for clim_p_ (see Appendix S2, Table S2 and Figure S2).

Linear mixed-effect models were fitted to quantify the effects of tree age, clim_p_ and clim_t_ on tree height. The model equation takes the form (Eq. 1):

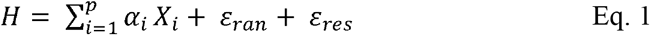

where *H* is tree height, *α_i_* is the set of *p* parameters associated with the main and interactive fixed effects of *X_i_* (tree age, clim_p_, clim_t_), *ε_ran_* is the variance component associated with the random terms, and *ε_res_* is the residual distributed error, usually following a Gaussian distribution (see Results section).

The saturated model for the fixed part, 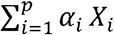, contained the linear and quadratic terms for each explanatory variable and all the potential pair-wise and three variable interactions (i.e. 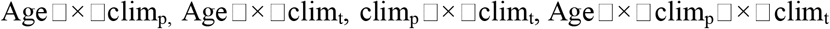). The random part of the model allowed us to consider three dimensions of common gardens experiments: a) the hierarchical nature of the data derived from the experimental design (i.e. populations nested within blocks, and blocks nested within trials), b) the temporal correlation between repeated measurements within tree individuals (i.e. individual tree), and c) the potential sources of variation not included in the fixed effects (such as soil, variation occurring at smaller spatial scales than blocks, etc.). All the variables were examined for outliers and departures from normality and the linearity of the relationships of each predictor with the response variable was checked (through residual plots for each predictor variable in the final model).

We selected the best-supported model starting from a saturated model following a hierarchical backward selection procedure (Burnham & Anderson, 2002; Zuur *et al*., 2009). For the random part of the model, we selected the structure with the lowest AIC value (round 1). For the fixed part, we used the Akaike Information Criterion (AIC) (Akaike, 1992), following the rule that net increments of lower than two units of AIC associated with the elimination of any parameter in the full model determined the exclusion of the parameter from the final model. We started by testing the three-variable interaction (round 2), followed by the two-variable interaction (round 3), main effects (round 4), and linear effects (round 5).

Differences in AIC between models allowed us quantifying the relative importance of each predictor variable. The random effects were tested using restricted maximum likelihood of the parameter (REML), and fixed effects using maximum-likelihood (ML). Finally, parameter estimates of the best-supported model were obtained using restricted maximum likelihood (REML), which minimizes the likelihood of the residuals from the fixed-effect portions of the model (Zuur *et al*., 2009). The variance explained by the model was assessed by pseudo-R^2^ (Nakagawa & Schielzeth, 2013) that splits the variance into the marginal MR^2^ (explained solely by the fixed effects) and conditional CR^2^ (explained by fixed and random effects together). This pseudo-R^2^ cannot be calculated with all the combinations between the family distributions and link functions (e.g. Gaussian family with identity link “log”) for linear mixed-effect models, therefore the goodness of fit of the models was also assessed by computing the capacity of generalization of the model (CG). To do this, we calculated the Pearson coefficient, ρ, between a model fitted with the 2/3 parts of the data and independently validated with the remaining 1/3 part of the data. Additionally, to detect collinearity between explanatory variables in the best-supported model we used the variance inflation factor (VIFs) and set maximum value of VIF to 5 which is considered acceptable (Belsley, 1991). Appropriateness of the best model was assessed by plots of predicted vs. observed values. We used the R version 3.2.3 (R Core Team, 2016) run in linux-gnu operating system to perform all the analyses, and the ‘‘lme4’’ package (Bates *et al*., 2015).

#### 2. Computing populations’ phenotypic responses

Using the best-supported model for each species, we predicted populations’ phenotypic responses curves of tree height across the climatic range covered by the trials, clim_t_, for the three developmental classes (DC). Specifically, we fixed tree age using the DC, and the climate of origin of each population (clim_p_), and then we predict tree height responses curves along the climate of the trial (clim_t_) varying between the 99% percentiles observed in clim_t_ data.

#### 3. Computing phenotypic plasticity indices

Using the populations’ phenotypic responses curves of tree height and developmental classes (DC.1, DC.2 and DC.3), we computed two phenotypic plasticity indices across the climatic ranges covered by the trials (reviewed in Valladares *et al*., (2006)).

1) Phenotypic plasticity index (PP) computed as follows:

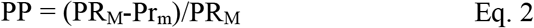

where PR_M_ is the highest phenotypic value across the population’s phenotypic response and across the climatic range studied. PR_m_ is the lowest phenotypic value across the climatic range studied. This index ranges between values of zero and one. The closer the values are to zero the less plastic the population is; and the opposite with values close to one.

2) Coefficient of variation of the phenotypic response (CV) computed as follow:

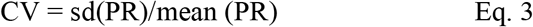

where PR is the phenotypic value at each point of the climatic range studied across the population’s phenotypic response fitted. sd is the standard deviation. This index reflects well the range of phenotype variation across the studied range. This index ranges between zero and one; the smaller the CV is, the smaller the plasticity is; and the opposite, the greater the CV is, the greater the plasticity is.

#### 4. The developmental class effect on populations’ tree height plasticity indices

We tested if plasticity in tree height changed with the developmental class within species. To this aim, we performed analyses of variance of the two phenotypic plasticity indices, PP and CV, for the developmental classes and post-hoc pairwise comparisons of Tukey HSD (Honestly Significant Difference).

#### 5. The inter-annual climate variation during the 20^th^ century effect on tree height plasticity indices

For each species and each developmental class, we tested if inter-annual climate variation at the populations’ origin could explain the current degree of phenotypic plasticity measured by the two phenotypic plasticity indices (PP and CV). To this end, we fitted linear fixed-effect models between the phenotypic plasticity index (PP or CV, as the response variable) and the set of inter-annual climate variation indices (sd bio1, sd bio2, sd bio5, sd bio6, sd bio12, sd bio13 and sd bio14, as explanatory variables) for each developmental class (Eq. 4). Collinearity in the models was controlled by including climate variation indices whose co-variation measured with the Pearson’s correlation coefficient was below |0.7|.

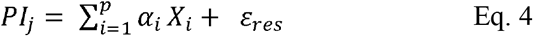

where PI_j_ is the phenotypic plasticity index at the developmental class j (j= 1, 2 or 3), *α_i_* is the set of *p* parameters associated with the effects of *X_i_* (sd bio_i_) and *ε_res_* is the residual error. Models were fitted with a Gaussian distribution of errors and identity link function. A step-wise procedure (direction= “backward”) was implemented to choose the best-supported model. Appropriateness of the models were assessed by plots of residuals vs. fitted values, qq-plots and the Cook’s distance that identify outliers in the data that could over-influence the model fitting, if necessary they were removed from the analysis.

## Results

### The model: Tree height responses accounting for tree age and climate

The model that included all the factors tested (tree age, clim_p_ and, clim_t_) with the linear and quadratic effects, and three and two-pairwise interactions, was the best-supported one for the three species (Table 1). The final model for *P. nigra* included age, mean annual temperature at the trial (bio1_t_) and annual water availability at the populations’ origin (WAI_p_); *P. pinaster* included age, annual potential evapotranspiration at the trial (PET_t_) and winter precipitation at the populations’ origin (prec.djf_p_); and for *P. pinea* included age, maximum temperature of the warmest month at the trial (bio5_t_) and summer precipitation at the populations’ origin (prec.jja_p_) (Table 2, Table S3 and Fig. S3). All models produced unbiased estimates of tree height and high capacity of generalization, as well as high marginal and conditional explained variance, CG/MR^2^/CR^2^, with 0.79/not available/not available, 0.72/0.83/0.96 and 0.80/0.69/0.97) for *P. nigra*, *P. pinaster* and *P. pinea,* respectively (Table 2).

**Table 1.**
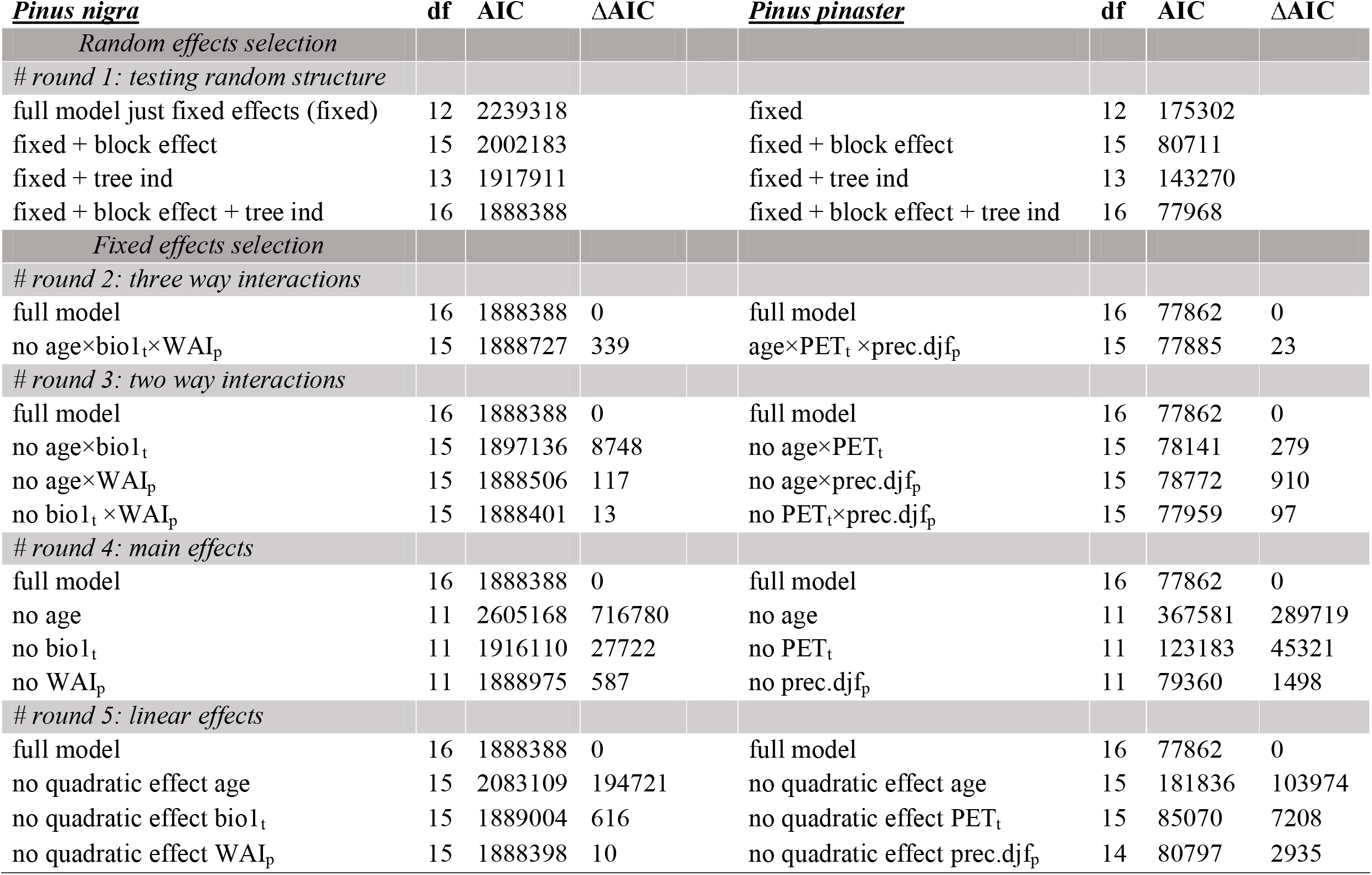
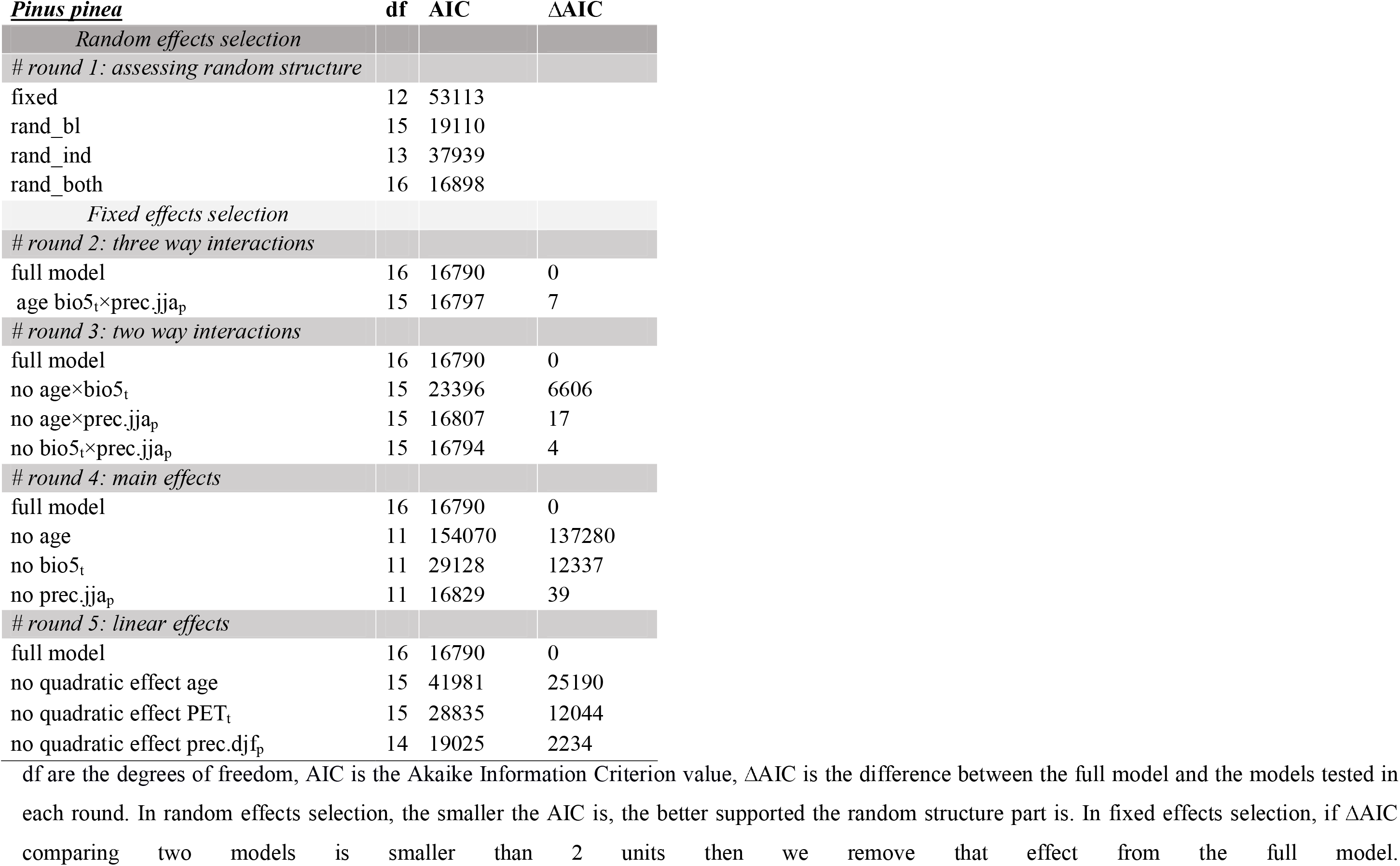
Random and fixed effects selection of tree height model in response to age, clim_t_ and clim_p_ using the Akaike Information Criterion (AIC) for the three pine species.

**Table 2.**
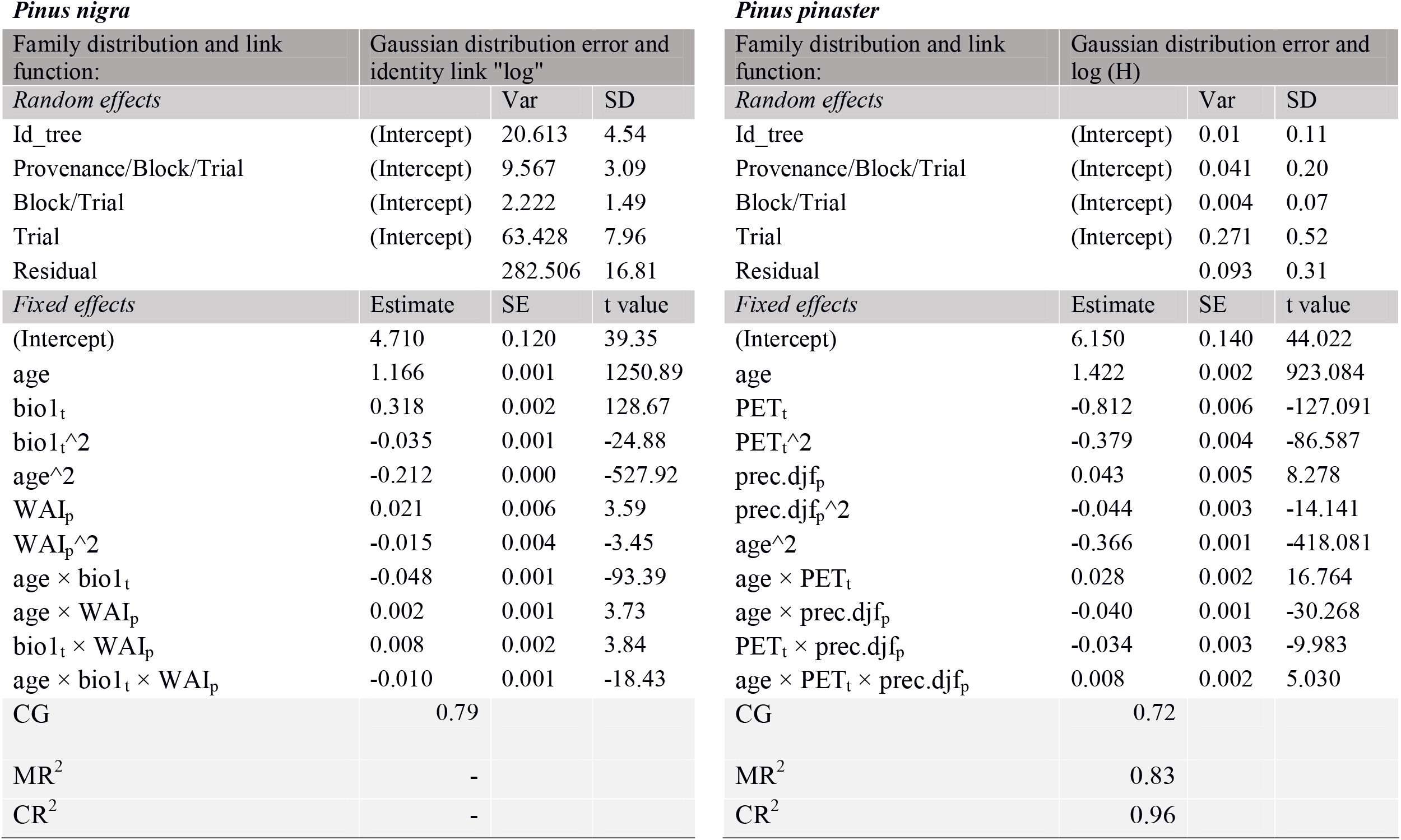
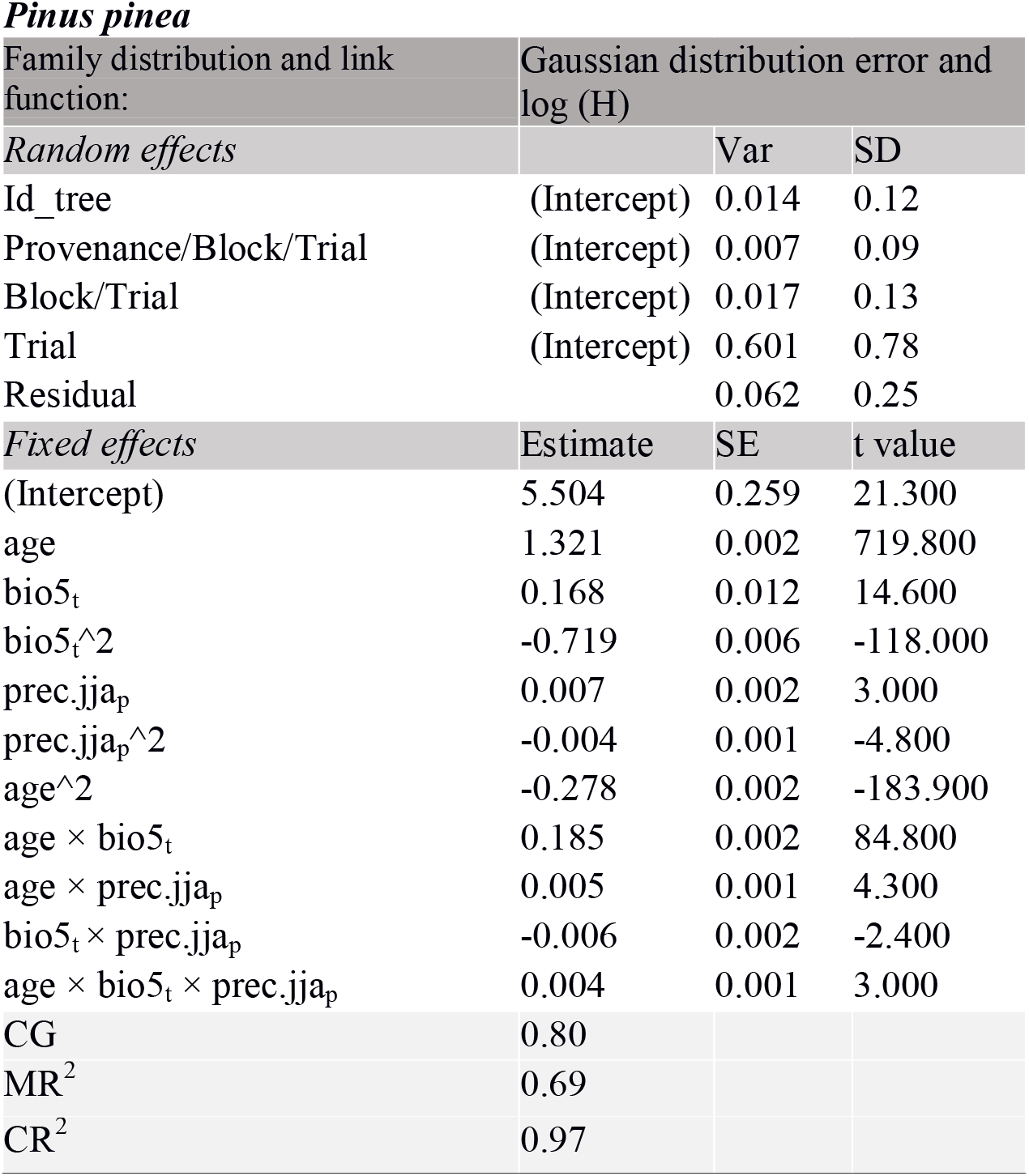
Detailed description of the best-supported model for each pine species analyzed including the family distribution and link function, the variance and standard deviation for the random effects (Var and SD respectively); and the estimated parameter, standard error and t-values for the fixed-effects (Estimate, SE and t-values, respectively).

### Main drivers of tree height triggering populations’ phenotypic responses

Overall, tree age made the largest contribution to tree height, followed in order of importance by the climate at the trial and at the populations’ origin: clim_t_ and clim_p_, respectively (Table 1; see ΔAIC comparisons). The mean annual temperature of the trial presented a positive effect on tree height in *P. nigra* (Fig.1a and Fig.S4a), but in the other two species, at a certain evaporative demand (either expressed in mm by annual potential evapotranspiration or degrees Celsius by the maximum temperature of the warmest month), the temperature had a negative effect on tree height, see populations’ phenotypic responses in Fig.1 and Fig.S4 for *P. nigra* and *P. pinaster*.). The most important interaction was age with clim_t_, except in *P. pinaster* that was age with clim_p_ (Tables 1 & 2). Finally, the interaction term between clim_t_ × clim_p_ overall contributed the least to tree height, although in *P. pinaster* this contribution was higher than in the other species (Table 1 and populations’ phenotypic responses in Fig.1. and Fig.S4.).

**Figure 1.**
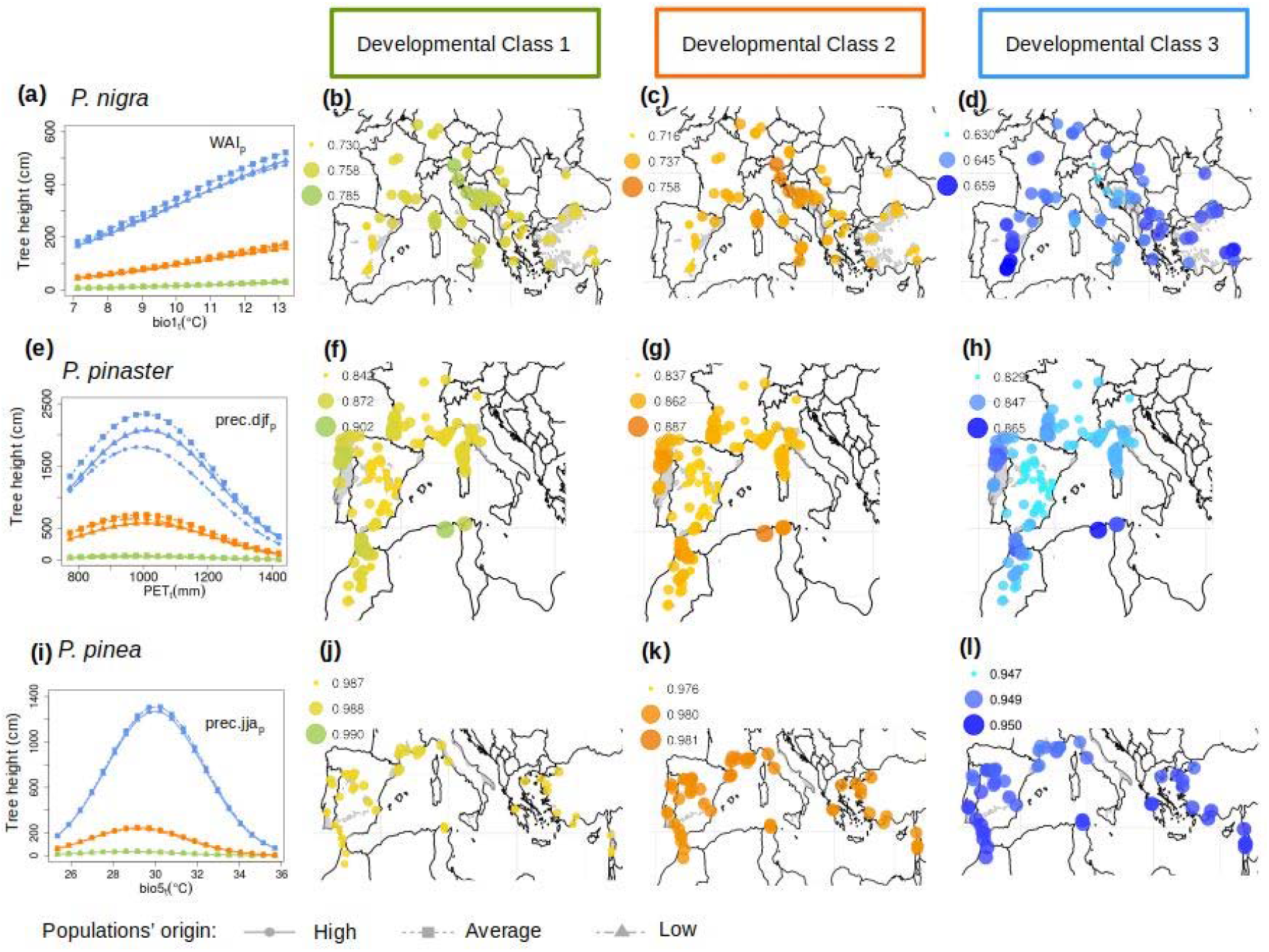
Populations’ phenotypic tree height responses across clim_t_ particularized for three populations’ origin (High, Average and Low in terms of clim_p_ values) and for the three developmental classes, DC, (Developmental Class 1: green, DC. 2: orange and DC. 3: blue) for a) *P. nigra*, e) *P. pinaster* and i) *P. pinea*. Values of the phenotypic plasticity index (PP) for the three developmental classes across the species natural distribution ranges are shown. DC. 1: b), f) and j), DC. 2: c), g) and k); and DC. 3: d), h) and i).

### The developmental class effect on populations’ tree height plasticity indices

Overall, phenotypic plasticity indices decreased significantly across developmental classes, i.e. young trees are the most plastic ones (Fig. 2, Tables S4 and S5). The greatest values of plasticity were found for *P. pinea* and the least for *P. nigra* for all developmental classes (Fig. 2). Finally, intraspecific phenotypic plasticity variation was the greatest in *P. pinaster* and the lowest in *P. pinea* (Fig. 2).

**Figure 2.**
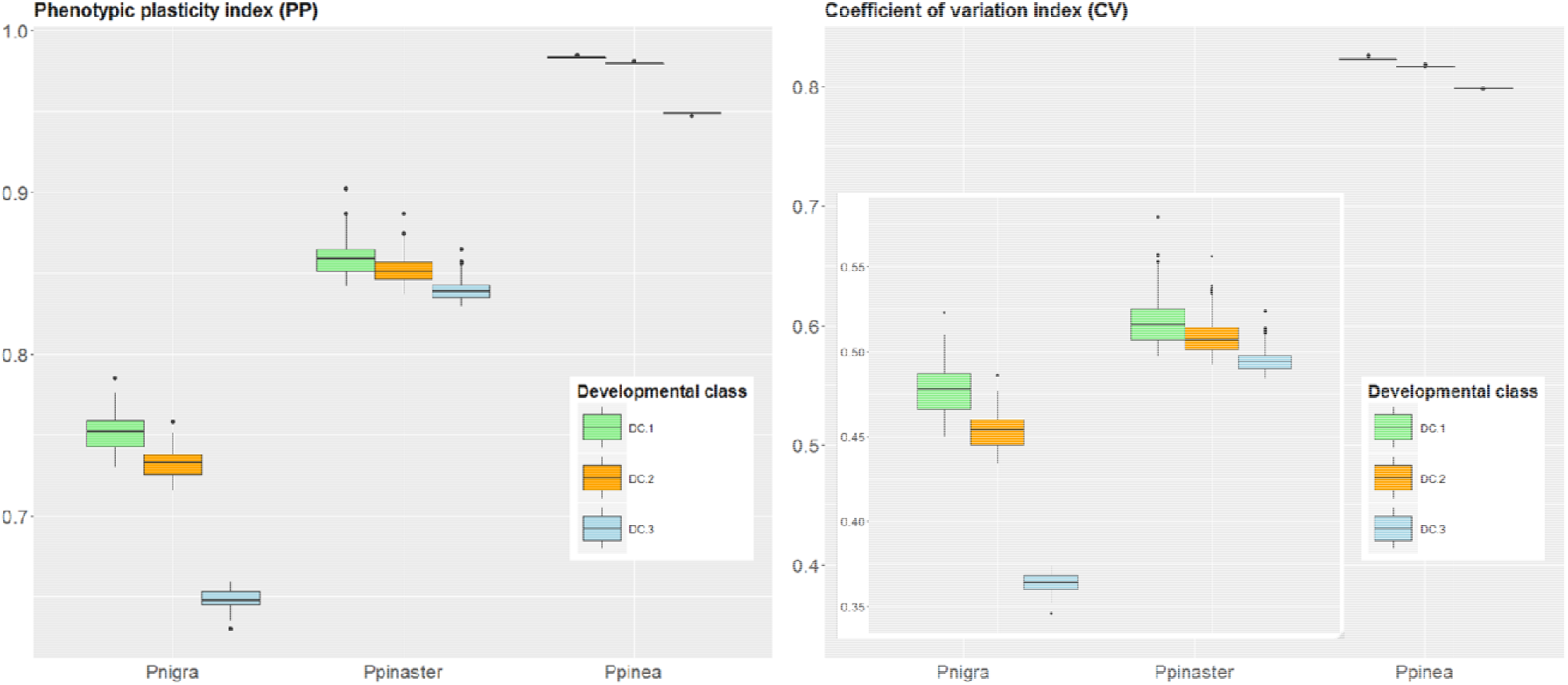
Phenotypic plasticity values for the three species and for the three developmental classes, DC, (Developmental Class 1: green, DC. 2: orange and DC. 3: blue) for the two indices computed (PP and CV). An inset graph is included in the CV index as the values of *P. nigra* and *P. pinaster* are significantly smaller compared with those obtained in *P. pinea*.

### The inter-annual climate variation during the 20^th^ century effect on tree height plasticity indices

For the three species, we did not include the standard deviation of annual precipitation (sd bio12) in the fixed-effect models because it was highly correlated with the standard deviation of the precipitation of the wettest month (sd bio13): Pearson’ correlation coefficients of 0.76, 0.82 and 0.91 for *P.* nigra, *P. pinaster* and *P. pinea*, respectively. Moreover, we removed some populations whose Cook’s distances were above 1 and over-influenced the fitted models (Appendix S3). Also, the model fitted for early adults in *P. pinea* using the CV index was not included in the results as it did not accomplished the linearity assumptions.

Overall, inter-annual temperature and precipitation variation during the 20^th^ century in the standard deviation (sd) of the maximum temperature of the warmest month (sd bio5), sd of the precipitation of the wettest month (sd bio13) and sd of the precipitation of the driest month (sd bio14) were positively correlated with phenotypic plasticity indices (Table 3); while inter-annual variation in sd of the average annual temperature (sd bio1) was negatively correlated with phenotypic plasticity indices, with the exception of the developmental class 3 in *P. nigra* and *P. pinea* (Table 3). The variance explained by the phenotypic plasticity indices models (P.I.) was high, and ranged between 0.69 and 0.70 for *P. nigra*, 0.80 and 0.84 for *P. pinaster*, and 0.71 and 0.76 for *P. pinea* (Table 3).

**Table 3.**
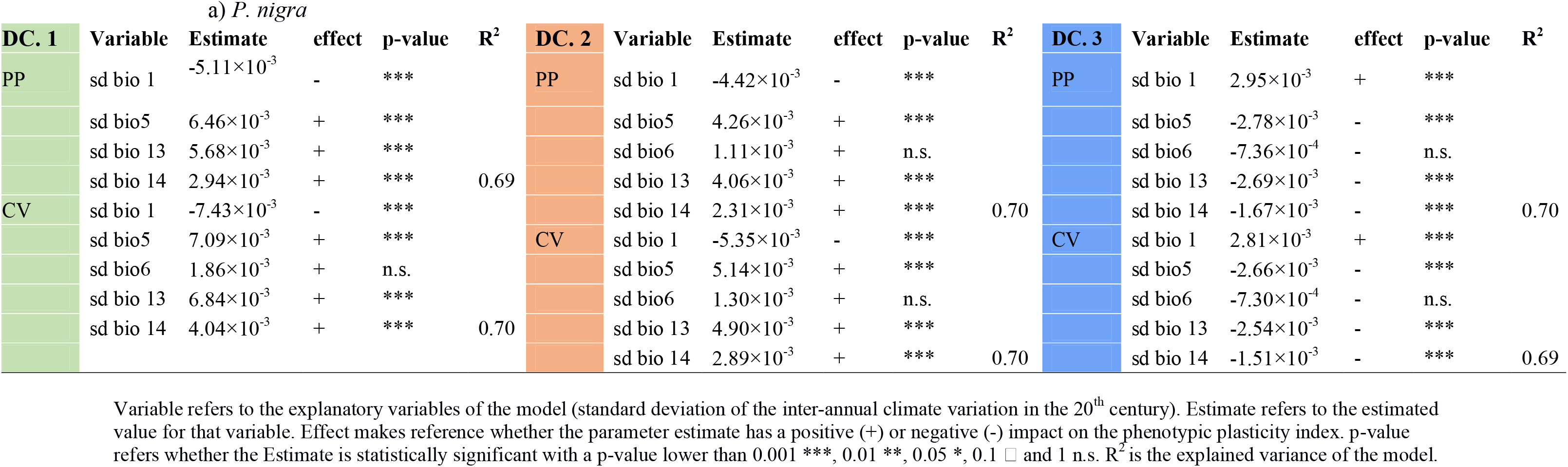
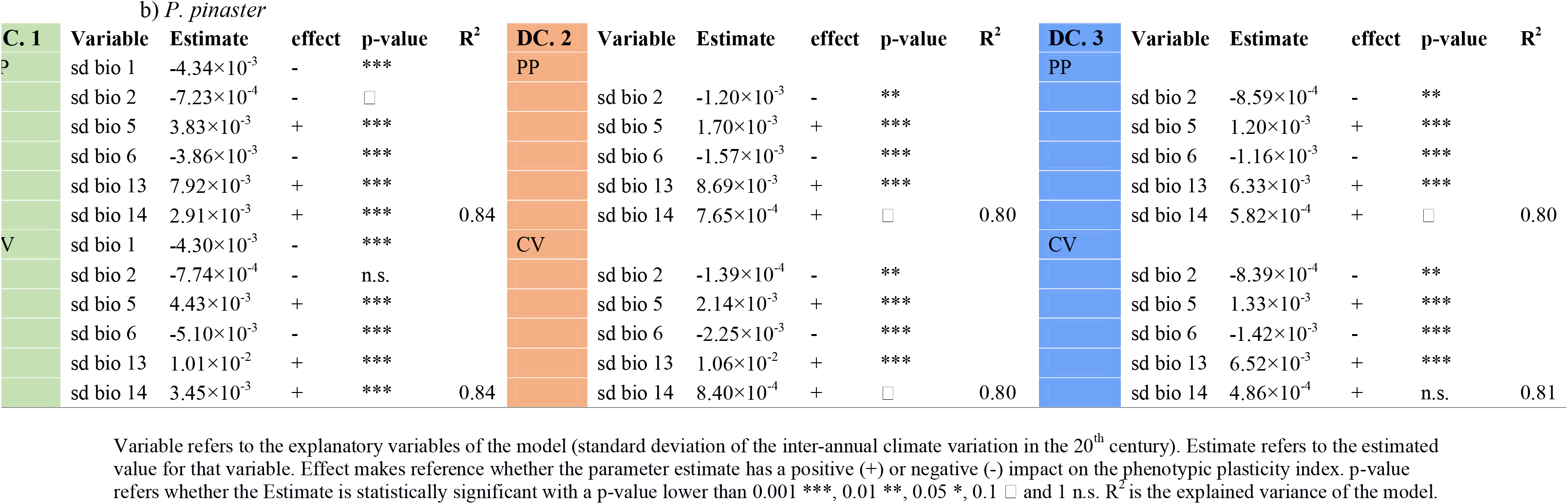
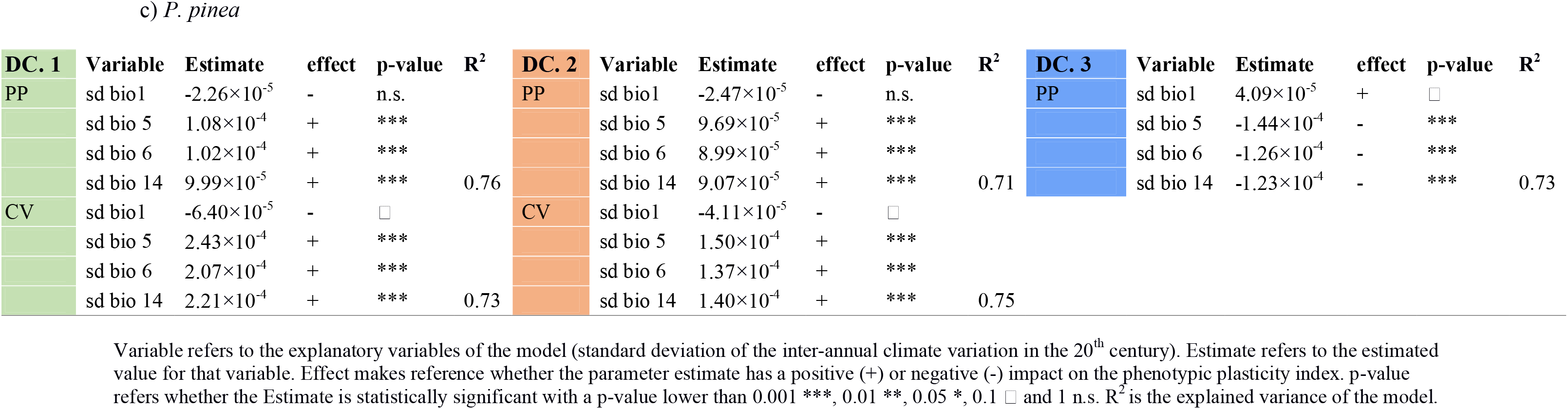
Results from the linear fixed-effect models between phenotypic plasticity indices (P.I.) and the standard deviation climate variables (sd bio). Three sub-tables are presented for each pine species, a), b) and c). Each sub-table shows the results for the two indices and the three developmental classes (DC) analyzed. Developmental Class 1: green, DC. 2: orange and DC. 3: blue

The results of the two indices were similar (Table 3). For illustrative purposes we plot the PP and CV indices along the sd bio5 variable that was statistically significant in all the models tested. Overall, we found that populations that experienced higher inter-annual climate variation during the 20^th^ century (sd bio5) presented higher plasticity in tree height for the three developmental classes, except in the developmental class 3 for *P. nigra* and *P. pinea* (Fig.3 and Fig.S5). Spatial differences among populations of PP and CV were also similar, with more contrasted differences among populations for *P. pinaster* (Fig. 1 and Fig. S5).

**Figure 3.**
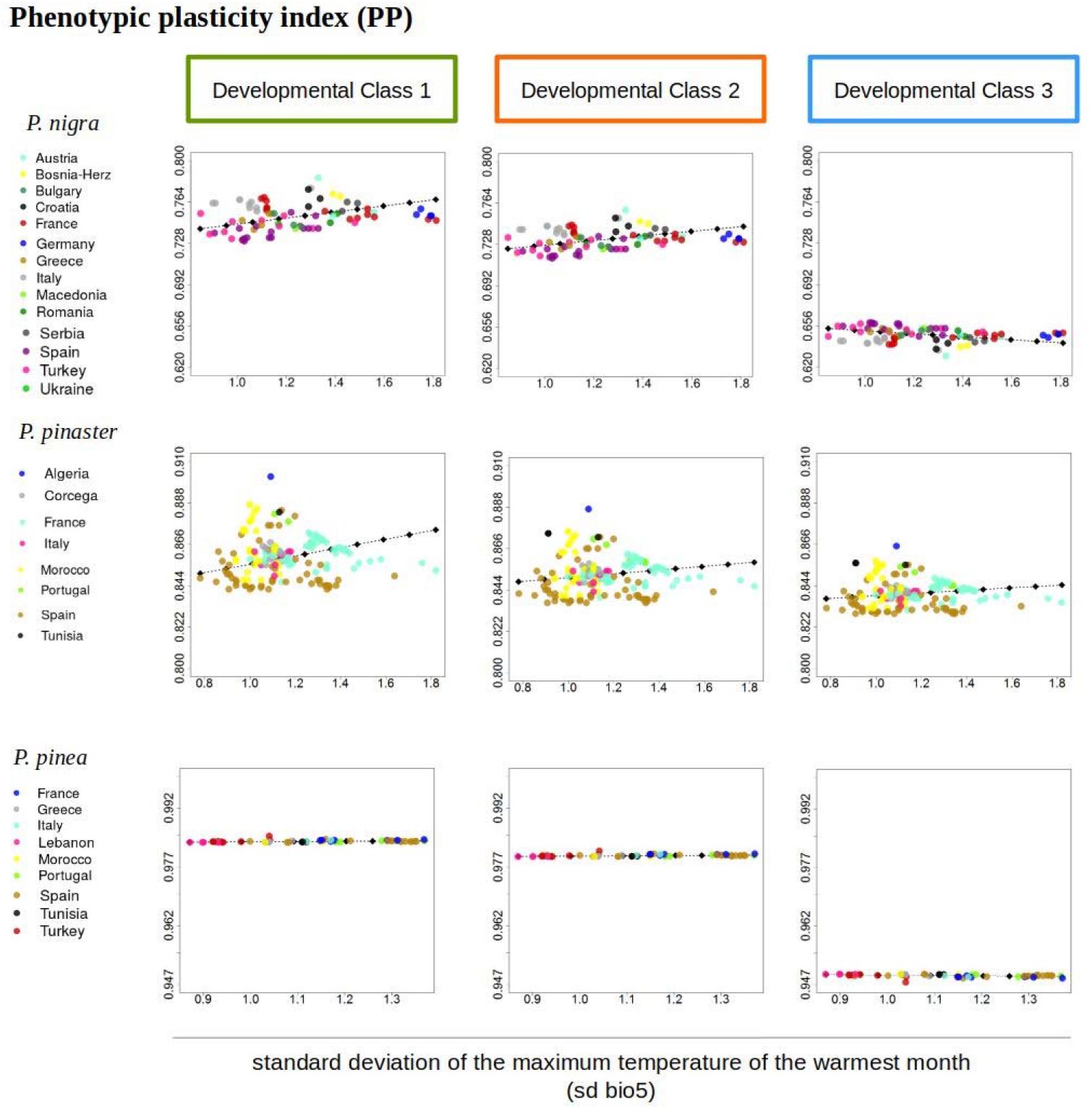
Phenotypic plasticity predictions of the PP index across the standard deviation of the maximum temperature of the warmest month (sd bio5) for the three developmental classes and pines species.

## Discussion

The use of range-wide multi-year tree height measurements compiled on common gardens allowed us to quantify the effect of tree age, population differentiation and plasticity across the distribution ranges of three Mediterranean pines. Overall, our results show that: (i) A significant part of phenotypic plasticity in tree height was explained by the inter-annual climate variation during the 20^th^ century under which tree populations evolved; (ii) Younger trees were more plastic in tree height than older trees; (iii) Although populations’ responses to climate were largely driven by phenotypic plasticity we did not find a geographical pattern of phenotypic plasticity across the species’ ranges.

### Main climatic drivers of tree height triggering populations’ phenotypic responses

Plastic responses in tree height were mainly driven by temperature-related variables (annual mean temperature, annual potential evapotranspiration and maximum temperature of the warmest month), and in general, rising temperatures led to higher trees up to a certain value. This result is in agreement with previous studies suggesting that higher heights in pines are found at warmer sites, if drought is not limiting (Vizcaíno-Palomar *et al*., 2016). This could be explained because warm temperatures, up to a certain threshold, allow trees to have higher photosynthetic capacity, resulting in a higher rate of carbon assimilation (Way & Oren, 2010) but going beyond that threshold, it can imply the opposite effect. Accordingly, in *P. pinaster* and *P. pinea* tree height decreases when the evaporative demand is too high due to stomatal closure and reduction of the photosynthetic activity (Pasho *et al*., 2012; Mazza *et al*., 2014). However, we did not find that threshold in *P. nigra.* This species is found at higher altitudes than *P. pinaster* and *P. pinea*, in mountainous areas where high temperatures can be counterbalanced by the altitude effect and hence allow for increments of tree height growth. Moreover, it can be explained by the fact that the range of climate covered by the trials do not cover the complete population’s phenotypic response, explaining the lack of a maximum tree height as it have been found in the other two species. Altogether, these results suggest that high temperatures linked with water stress are the main climatic drivers liming tree height in the three pine species studied.

The main driver of population differentiation (population effect) in tree height was precipitation (for *P. pinaster* and *P. pinea*) and annual water availability (for *P. nigra*). This points out to the selective role of water availability across the distribution range of these mostly Mediterranean trees (Pigott & Pigott, 1993). Our findings suggest that evolutionary processes in tree height were mostly driven by water availability (Aranda *et al*., 2009; Sánchez-Gómez *et al*., 2011), although local adaptation is driven by minimum winter temperatures for *P. nigra* (Kreyling *et al*., 2012), and by mean annual temperature for *Pinus pinea* (Mutke *et al*., 2010). The highest differences in tree height among populations were found for *P. nigra* and *P. pinaster* (Fig. 1a. and Fig. 1e.). In general, populations originating from the extremes of the climatic gradient, either under high or low values in rainfall or water availability, underperformed compared to populations originating from intermediate climates, but these differences are more marked in *P. pinaster.* For example, *P. pinaster* populations from the south of the distribution are better adapted to drought: they invest higher biomass to root and less to stem development than populations from northern parts of the distribution (Aranda *et al*., 2009). In *P. nigra*, differences in tree height due to genetic effects have also been recorded (Thiel *et al*., 2012), whereas *P. pinea* shows low genetic variation among populations (Fig.1i.). This is in agreement with previous studies reporting little genetic variation in morphological and physiological quantitative traits -e.g. photosynthesis, biomass partitioning, SLA, etc. (Court-Picon *et al*., 2004; Mutke *et al*., 2010; Sánchez-Gómez *et al*., 2011) but null in Chambel *et al*., (2007).

### The developmental class effect on populations’ tree height plasticity indices

Young pine trees were more plastic than early adults (Fig. 2). This result suggests that the capacity to respond plastically changes along the life cycle of trees. Phenotypic plasticity differ among species, among populations, among traits (Valladares *et al*., 2002; Bradshaw, 2006), and here we show that it also varies with age. The first stages of recruitment are critical for plant establishment, and hence greater capacity of plasticity in tree height can be advantageous to avoid competition and reach light. In addition, small changes in the environment can be more noticeable for seedlings than to saplings or adult trees that are already well established with their root systems installed into deeper layers of the soil compared to seedlings. For instance, soil moisture variation is higher in the shallow layers of the soil than in deeper ones, hence promoting greater phenotypic plasticity. As a consequence, phenotypic plasticity variation across developmental classes can impact into many ecological processes, such as population and community dynamics, the community assembly and ecosystem functioning.

### The inter-annual climate variation during the 20^th^ century effect on tree height plasticity indices

Our findings suggest that plasticity in tree height is a trait that is under selection driven by climate variability (Table 3, Fig. 3 and Fig. S5). This finding is consistent for the three Mediterranean species regardless the origin of the populations. Populations that evolved under high inter-annual climate variation, in either maximum or minimum values in temperature or precipitation of climate variables associated with extreme values (standard deviation of the maximum temperature of the warmest month, sd of the precipitation of the wettest month and sd of the precipitation of the driest month) during the 20^th^ century, have great capacity to respond plastically in tree height to changes in climate (Fig. 3), regardless their position at the core or at the margin of the distribution range (maps in Fig. 1 and Fig.S4). These results are in agreement with previous studies in plant species where plastic responses were associated with climate variation in *Convolvulus chilensis* and *Senna candolleana* (Gianoli & González-Teuber, 2005; Lázaro-Nogal *et al*., 2015). Although local adaptation (population effect in common garden data) seems to clearly follow a geographical pattern in European trees (including the ones studied here) (Frejaville *et al*.), our results show that plasticity geographical patterns are more complex (Valladares *et al*., 2014).

### Implications of phenotypic plasticity for evolutionary responses to climate change

Our findings are important in the context of climate change because plastic genotypes would likely increase their odds to persist at the short-time if plasticity is adaptive, and can also be advantageous if plastic genotypes are subject to further evolution that promotes the necessary genetic changes to reach the new optimum and get adapted to the new environment (Pigliucci, 2005; Richards *et al*., 2006). Among the three studied species, *P. pinaster* presents high values of plasticity in tree height combined with a high differentiation among populations (Fig. 1e. and Fig. 2), suggesting good chances to respond to climate change in the short term by phenotypic plasticity and keeping evolutionary potential to adapt in the long term. *P. pinea* presents the highest phenotypic plasticity values out of the three studied species, but combined with low differentiation among populations and low genetic diversity (Fig. 1i. and Fig. 2), which makes plasticity virtually the unique way for this species to respond to changes in the environment. However, we cannot rule out that plasticity for tree height is related to higher fitness and our results call into question whether higher plasticity could be adaptive and hence beneficial to cope with climate change in the short-term.

## Limitations

Although the network of provenance tests used in the present study cover relatively well the species distribution ranges, our results of phenotypic plasticity in tree height are confined to the climatic gradients covered by these common gardens. For example, in *P. nigra* phenotypic plasticity values could have been underestimated because the maximum tree height is not reached within the mean annual temperature range studied. Improved results regarding phenotypic plasticity could be obtained by establishing new common gardens both within the species’ distribution to complete the range and outside the climatic range of the species.

## Conclusions

Under a climate change context, the potential of the three species to persist *in-situ* largely rely on their plastic responses, regardless of their genetic diversity. The predominance of the plastic effect over the genetic one highlights that at the short-term, species’ strategy to keep pace with climate change will likely rely on eco-physiological adjustments to environmental changes rather than on evolutionary responses. However, our current understanding of plasticity makes difficult to ascertain if plasticity will be adaptive and in case, which will be the real limits of plasticity. Therefore, to allow species persistence in the long-term, genetic variation within populations is essential to respond by evolutionary processes to environmental changes. Likewise, our results call into question whether future climate change variation would promote plasticity in the near future as our results showed that happened in the 20^th^ century.

## Data Accessibility Statement

All phenotypic data used in this study are available on ZENODO with DOIs 10.5281/zenodo.3250704, 10.5281/zenodo.3250698 and 10.5281/zenodo.3250707 for *P. nigra*, *P. pinaster* and *P. pinea*, respectively (Vizcaíno-Palomar *et al*., 2019).

## Appendix S1 Detailed description for climate variable selection

We used two analysis to select the short-term climate related to the trial (clim_t_) and the long-term climate related to the climate at the population origin (clim_p_): i) linear mixed-effect models and ii) principal component analyses.

### i) Linear Mixed-Effect Models

To select clim_t_, we ran 21 linear mixed-effects models to analyse the response of tree height to each of the 21 climate variables to the climate of the trials (Table S1). To select clim_p_, we ran 21 linear mixed-effects models to analyse the response of tree height to each of the 21 climate variables to the climate at the populations’ origin. Random effects included populations nested into blocks, and those nested within trials, and trees nested within population, block and trial, to control for differences among sites and populations, and to control for repeated measurements of the same trees, respectively. Fixed effects included tree age and the climate variable, including the linear and the quadratic forms, and the linear interaction between them. All climate variables were standardized (the mean was subtracted from each value and divided by the standard deviation). The model equation takes the form:

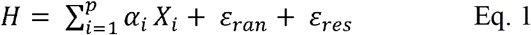

where *H* is tree height, *α_i_* is the set of *p* parameters associated with the main and interactive effects of *X_i_* climate (either clim_t_ or clim_p_) and age variables, *ε_ran_* is the variance component associated with the random terms, and *ε_res_* is the residual distributed error.

We selected the model in which the climate variable met three requisites: 1) the sign of the estimated quadratic coefficient must be negative in order to get concave responses (maximum tree height is expected at intermediate climatic values, decreasing towards the extremes) 2) high absolute values of the estimated coefficients as an approach of the size effect of the climate variable on tree height, 3) being biological meaningful variable. For example, potential evapotranspiration climate variable can provide us with more information than just a variable of annual precipitation. Usually, curve responses between clim_t_ and tree height are more likely to be concaves than tree height with clim_p_ (personal observation from preliminary analysis). Therefore for clim_t_ we focused on the absolute highest value of the negative estimated parameter in the quadratic term when statistically significant, while for clim_p_, we focused on whether the linear and quadratic terms were statistically significant and then we chose among those variables with the absolute highest negative parameter in the quadratic term. If only the linear parameter was significant, then we chose the variable with the highest estimated parameter.

### ii) Principal Component Analysis

We ran two independent principal component analyses (PCA) in R (R Core Team, 2016) for clim_t_ and clim_p_ variables for each species. These analyses help selecting the variable which is highly representative of the climate variation covered by the data. We generally choose the climate variable that belongs to the first PCA axis, which captures a higher variance of the data. However, sometimes we needed to choose the climate variable from the second PCA axis if the climate variable is highly correlated with the first PCA axis.

These two complementary analyses were defined to facilitate climate selection, but this was not always straightforward.

## Appendix S2 Description of the results from the linear mixed-effect models and principal component analyses to select the climate variables of clim_p_ and clim_t_

*P. nigra.* Mean annual temperature, bio1, was selected for clim_t_. It showed the highest size effect among the total of the climate variables tested in the linear mixed-effect models (Table S2). Moreover, mean annual temperature was highly correlated with the first PCA axis (0.76) and showed a moderate-high contribution to the first axis (6.37% being the maximum 9.36%) (Figure S2). Annual water availability, WAI, was selected for clim_p_. Although none of the tested climate variables in the linear mixed-effect models were statistically significant, the principal component analysis facilitated to choose it (Table S2). Thus, annual water availability was highly correlated with the first PCA axis (−0.93) and it showed the highest contribution to the first PCA axis, 8.33% (Figure S2).

*P. pinaster.* Annual potential evapotranspiration, PET, was selected for clim_t._ Although precipitation of the driest month could have been selected for modeling, we chose annual potential evapotranspiration as it was statistically significant in the linear mixed-effect models and it integrates temperature and precipitation values which make this variables more informative (Table S2). Annual potential evapotranspiration was highly correlated with the first PCA axis (0.94), and it showed the second highest contribution (7.57%) to this axis after summer daily mean temperature (Figure S2). Winter precipitation, prec.djf, was selected for clim_p_. It was statistically significant in the linear mixed-effect models, and the two estimated coefficients were biologically meaningful and their size effects were the highest (Table S2). The variable of winter precipitation was moderate-to-highly correlated with the second PCA axis (0.76), with a moderate-to-high contribution to the axis (8.21%, being the maximum contribution of 12.55%) (Figure S2). The second axis of the PCA explained nearly the same amount of variability, 33.38%, compared to the variance explained by the first axis of 45.19%.

*P. pinea.* Maximum temperature of the warmest month, bio5, was selected to represent clim_t_. It showed the highest size effect on the estimated quadratic term in the linear mixed-effect models. Moreover, maximum temperature of the warmest month was highly correlated with the first PCA axis (−0.91) and it showed a moderate-to-high high contribution to the first axis of the PCA (6.88%, being the maximum 7.86%) (Figure S2). Summer precipitation was selected to represent clim_p_. Both of the terms estimated, the linear and the quadratic terms, were statistically significant. The linear and quadratic terms were statistically significant. Moreover, summer precipitation was moderately correlated with the second PCA axis (−0.59) and it showed a moderate contribution to the axis (4.91%, being the maximum 11.51%) (Figure S2). The second axis of the PCA explained nearly the same amount of variability, 33.3% compared to 38.83%.

## Appendix S3 Populations’ with a Cook’s distance value above one

The Cook’s distance measures the effect of deleting a given population due to the presence of large residuals that can influence the accuracy of the model. We removed those populations from the linear-fixed effect models. Specifically, in *P. nigra* we deleted the population of Parapluberg in the three developmental classes (DC.1, DC. 2 and DC.3). In *P. pinaster*, we deleted four populations (the population of Val Freda in DC. 3, and the populations of Ain Baccouche, Tabarka and Valencia in DC. 1). In *P. pinea*, we deleted the population of Artvin in DC1, DC2 and DC3.

## Supporting Information

**Table S1.** Climatic variables used from the EuMedClim (Fréjaville & Benito Garzón, 2018).

| **Annual climate variables** |  |  |
| --- | --- | --- |
| **Number** | **Variable name** | **Description** |
| **1** | bio1 | annual daily mean temperature (° C) |
| **2** | bio2 | mean diurnal temperature range (max-min, ° C) |
| **3** | bio5 | maximum temperature of the warmest month, (° C) |
| **4** | bio6 | minimum temperature of the coldest month (° C) |
| **5** | bio12 | annual precipitation (mm) |
| **6** | bio13 | precipitation of the wettest month (mm) |
| **7** | bio14 | precipitation of the driest month (mm) |
| **8** | PET | annual potential evapotranspiration [PET] (mm) |
| **9** | WAI | water availability [bio12- PET] (mm) |
| **Seasonal climate variables** |  |  |
| **Number** | **Variable name** | **Description** |
| **10** | tmean.dfj | winter daily mean temperature (° C) |
| **11** | tmean.mam | spring daily mean temperature (° C) |
| **12** | tmean.jja | summer daily mean temperature (° C) |
| **13** | tmean.son | autumn daily mean temperature (° C) |
| **14** | prec.djf | winter precipitation (mm) |
| **15** | prec.mam | spring precipitation (mm) |
| **16** | prec.jja | summer precipitation (mm) |
| **17** | prec.son | autumn precipitation (mm) |
| **18** | PET.min | PET of the wettest month (mm) |
| **19** | PET.max | PET of the driest month (mm) |
| **20** | WAI.min | WAI of the driest month (mm) |
| **21** | WAI.max | WAI of the wettest month (mm) |

**Table S2.** Results from linear mixed-effect models to select the climate variables of clim_p_ and clim_t_. Bold letters indicate the selected variables. Complementary results from the Principal Component Analyses are shown in Figure S2

| *Pinus nigra* | Short term climate effects |  | Long term climate effects |  |  |  |  |
| --- | --- | --- | --- | --- | --- | --- | --- |
| Variable | coef<sup>2</sup> | t value | Variable | coef | t value | coef<sup>2</sup> | t value |
| **bio1<sub>t</sub>** | **-0.24** | **-112.12** | prec.jja<sub>p</sub> | 0.00 | -0.01 | -0.04 | -0.28 |
| tmean.djf<sub>t</sub> | -0.20 | -140.38 | PET<sub>p</sub> | -0.02 | -0.12 | -0.03 | -0.19 |
| PET.min<sub>t</sub> | -0.13 | -45.51 | WAI.min<sub>p</sub> | -0.02 | -0.08 | -0.02 | -0.14 |
| bio6<sub>t</sub> | -0.10 | -127.72 | bio5<sub>p</sub> | -0.02 | -0.11 | -0.02 | -0.18 |
| prec.jja<sub>t</sub> | -0.08 | -84.68 | PET.max<sub>p</sub> | -0.01 | -0.07 | -0.02 | -0.10 |
| prec.son<sub>t</sub> | -0.07 | -180.30 | **WAI<sub>p</sub>** | **-0.01** | **-0.03** | **-0.01** | **-0.08** |
| bio14<sub>t</sub> | -0.03 | -149.63 | bio2<sub>p</sub> | -0.02 | -0.09 | -0.01 | -0.06 |
| tmean.son<sub>t</sub> | -0.03 | -19.31 | prec.djf<sub>p</sub> | -0.05 | -0.20 | -0.01 | -0.08 |
| bio13<sub>t</sub> | -0.01 | -29.53 | tmean.jja<sub>p</sub> | -0.02 | -0.09 | -0.01 | -0.12 |
| pet.max<sub>t</sub> | -0.01 | -11.58 | prec.son<sub>p</sub> | 0.00 | 0.00 | -0.01 | -0.04 |
| WAI.min<sub>t</sub> | -0.01 | -9.42 | WAI.max<sub>p</sub> | -0.07 | -0.27 | -0.01 | -0.05 |
| tmean.jja<sub>t</sub> | 0.01 | 10.06 | bio13<sub>p</sub> | -0.05 | -0.19 | -0.01 | -0.05 |
| prec.mam<sub>t</sub> | 0.01 | 22.98 | tmean.son<sub>p</sub> | 0.01 | 0.04 | -0.01 | -0.06 |
| bio5<sub>t</sub> | 0.02 | 28.96 | bio1<sub>p</sub> | 0.01 | 0.07 | 0.00 | -0.05 |
| bio12<sub>t</sub> | 0.02 | 65.66 | tmean.djf<sub>p</sub> | 0.06 | 0.31 | 0.00 | -0.01 |
| WAI<sub>t</sub> | 0.03 | 72.89 | tmean.mam<sub>p</sub> | -0.01 | -0.05 | 0.00 | 0.00 |
| PET<sub>t</sub> | 0.04 | 28.05 | bio6<sub>p</sub> | 0.05 | 0.26 | 0.00 | 0.00 |
| prec.djf<sub>t</sub> | 0.05 | 100.84 | prec.mam<sub>p</sub> | -0.01 | -0.04 | 0.00 | 0.00 |
| tmean.mam<sub>t</sub> | 0.05 | 41.13 | bio12<sub>p</sub> | -0.05 | -0.20 | 0.01 | 0.03 |
| bio2<sub>t</sub> | 0.13 | 92.50 | PET.min<sub>p</sub> | 0.04 | 0.16 | 0.01 | 0.04 |
| WAI.max<sub>t</sub> | 0.13 | 211.40 | bio14<sub>p</sub> | -0.04 | -0.18 | 0.03 | 0.12 |

| *Pinus pinaster* | Short term climate effects |  | Long term climate effects |  |  |  |  |
| --- | --- | --- | --- | --- | --- | --- | --- |
| Variable | coef<sup>2</sup> | t value | Variable | coef | t value | coef<sup>2</sup> | t value |
| bio14<sub>t</sub> | -79.13 | -77.05 | **prec.djf<sub>p</sub>** | **92.89** | **5.83** | **-116.90** | **-15.43** |
| **PET<sub>t</sub>** | **-62.09** | **-37.09** | bio13<sub>p</sub> | 124.55 | 10.02 | -95.65 | -13.56 |
| WAI<sub>t</sub> | -40.50 | -34.15 | bio2<sub>p</sub> | 58.20 | 5.07 | -75.43 | -11.05 |
| WAI.max<sub>t</sub> | -28.87 | -52.10 | PET.max<sub>p</sub> | 101.64 | 7.73 | -84.34 | -9.88 |
| Variable | coef^2 | t value | Variable | coef | t value | coef^2 | t value |
| tmean.son<sub>t</sub> | -25.77 | -21.76 | prec.mam<sub>p</sub> | 75.40 | 6.56 | -37.47 | -6.85 |
| bio2<sub>t</sub> | -17.74 | -27.77 | prec.jja<sub>p</sub> | 163.64 | 14.18 | -42.90 | -6.05 |
| WAI.min<sub>t</sub> | -15.79 | -24.15 | bio6<sub>p</sub> | 5.58 | 0.47 | -36.23 | -5.04 |
| bio5<sub>t</sub> | -11.70 | -6.69 | WAI.max<sub>p</sub> | 83.98 | 7.58 | -26.93 | -3.66 |
| bio12<sub>t</sub> | -7.56 | -9.46 | prec.son<sub>p</sub> | 35.45 | 3.26 | -33.29 | -3.21 |
| prec.djf<sub>t</sub> | -4.04 | -14.02 | bio14<sub>p</sub> | 123.28 | 8.70 | -14.10 | -1.89 |
| PET.min<sub>t</sub> | 3.78 | 1.71 | WAI<sub>p</sub> | 52.54 | 5.85 | -12.40 | -1.55 |
| bio13<sub>t</sub> | 5.18 | 15.19 | bio12<sub>p</sub> | 54.79 | 5.45 | -10.01 | -1.25 |
| bio6<sub>t</sub> | 10.16 | 13.75 | PET<sub>p</sub> | -39.41 | -4.07 | -6.07 | -0.97 |
| prec.son<sub>t</sub> | 11.66 | 49.06 | tmean.son<sub>p</sub> | 22.65 | 1.78 | -3.60 | -0.47 |
| PET.max<sub>t</sub> | 12.06 | 21.90 | PET.min<sub>p</sub> | -70.27 | -5.89 | -0.67 | -0.11 |
| tmean.djf<sub>t</sub> | 13.18 | 10.17 | tmean.djf<sub>p</sub> | 29.28 | 2.29 | -0.40 | -0.05 |
| prec.mam<sub>t</sub> | 14.62 | 27.30 | WAI.min<sub>p</sub> | 64.09 | 7.41 | 2.43 | 0.32 |
| prec.jja<sub>t</sub> | 18.26 | 15.69 | bio5<sub>p</sub> | 321.98 | 15.39 | 28.08 | 2.16 |
| tmean.jja<sub>t</sub> | 62.56 | 31.59 | bio1<sub>p</sub> | 49.67 | 3.57 | 21.93 | 2.87 |
| tmean.mam<sub>t</sub> | 89.81 | 71.02 | tmean.mam<sub>p</sub> | 63.97 | 4.65 | 24.23 | 3.20 |
| bio1<sub>t</sub> | 142.09 | 56.23 | tmean.jja<sub>p</sub> | 333.48 | 13.47 | 148.08 | 9.88 |

| *Pinus pinea* | Short term climate effects |  | Long term climate effects |  |  |  |  |
| --- | --- | --- | --- | --- | --- | --- | --- |
| Variable | coef^2 | t value | Variable | coef | t value | coef^2 | t value |
| **bio5<sub>t</sub>** | **-181.07** | **-152.89** | pet.mean<sub>p</sub> | 0.38 | 0.21 | -2.23 | -2.43 |
| prec.son<sub>t</sub> | -99.92 | -76.75 | bio5<sub>p</sub> | -1.61 | -0.80 | -2.07 | -1.91 |
| pet.min<sub>t</sub> | -54.20 | -29.40 | **prec.jja<sub>p</sub>** | **4.67** | **1.96** | **-1.88** | **-2.39** |
| tmean.jja<sub>t</sub> | -49.02 | -67.42 | PET.max<sub>p</sub> | 0.68 | 0.33 | -1.34 | -0.96 |
| bio1<sub>t</sub> | -21.73 | -24.04 | WAI.min<sub>p</sub> | -0.43 | -0.20 | -1.33 | -1.03 |
| bio6<sub>t</sub> | -17.76 | -35.58 | bio14<sub>p</sub> | 2.99 | 1.09 | -1.14 | -1.64 |
| bio14<sub>t</sub> | -11.35 | -26.11 | tmean.jja<sub>p</sub> | -4.86 | -2.47 | -1.08 | -0.75 |
| tmean.son<sub>t</sub> | -9.99 | -16.22 | tmean.mam<sub>p</sub> | -5.34 | -2.91 | -0.66 | -0.46 |
| tmean.mam<sub>t</sub> | -9.29 | -16.19 | tmean.djf<sub>p</sub> | -5.26 | -2.87 | -0.36 | -0.19 |
| PET.max<sub>t</sub> | -7.60 | -12.53 | PET.min<sub>p</sub> | -2.05 | -0.97 | -0.34 | -0.44 |
| WAI.min<sub>t</sub> | 2.66 | 9.36 | bio6<sub>p</sub> | -5.23 | -2.68 | 0.21 | 0.10 |
| prec.mam<sub>t</sub> | 5.11 | 13.96 | bio2<sub>p</sub> | 3.25 | 1.52 | 0.29 | 0.16 |
| bio13<sub>t</sub> | 7.00 | 10.69 | bio1<sub>p</sub> | -5.87 | -3.20 | 0.54 | 0.31 |
| prec.djf<sub>t</sub> | 12.97 | 45.43 | WAI<sub>p</sub> | -5.80 | -2.90 | 1.80 | 1.28 |
| WAI.max<sub>t</sub> | 22.84 | 38.69 | prec.son<sub>p</sub> | -9.19 | -3.92 | 2.59 | 2.26 |
| Variable | coef^2 | t value | Variable | coef | t value | coef^2 | t value |
| prec.jja<sub>t</sub> | 25.26 | 84.43 | tmean.son<sub>p</sub> | -7.19 | -4.05 | 3.71 | 2.13 |
| tmean.djf<sub>t</sub> | 35.07 | 47.88 | prec.mam<sub>p</sub> | -7.98 | -3.36 | 3.84 | 2.59 |
| bio2<sub>t</sub> | 38.08 | 20.62 | bio12<sub>p</sub> | -9.18 | -4.64 | 4.60 | 3.16 |
| PET<sub>t</sub> | 57.09 | 26.45 | bio13<sub>p</sub> | -13.95 | -6.45 | 8.76 | 5.41 |
| bio12<sub>t</sub> | 74.68 | 88.44 | ppet.max<sub>p</sub> | -14.19 | -6.45 | 9.41 | 5.28 |
| WAI<sub>t</sub> | 81.80 | 66.39 | prec.djf<sub>p</sub> | -15.37 | -6.54 | 9.61 | 5.70 |

**Table S3.** Variance inflator factors (VIF) of the best-supported model for each pine species analyzed.

| *Pinus nigra* |  | *Pinus pinaster* |  | *Pinus pinea* |  |
| --- | --- | --- | --- | --- | --- |
| Variable | VIF | Variable | VIF | Variable | VIF |
| age | 5 | age | 2 | age | 2 |
| bio1<sub>t</sub> | 1 | PET<sub>t</sub> | 2 | bio5<sub>t</sub> | 2 |
| bio1<sub>t</sub><sup>2</sup> | 2 | PET<sub>t</sub><sup>2</sup> | 2 | bio5<sub>t</sub><sup>2</sup> | 4 |
| age<sup>2</sup> | 5 | prec.djf<sub>p</sub> | 2 | prec.jja<sub>p</sub> | 2 |
| WAI<sub>p</sub> | 1 | prec.djf<sub>p</sub><sup>2</sup> | 2 | prec.jja<sub>p</sub><sup>2</sup> | 2 |
| WAI<sub>p</sub><sup>2</sup> | 1 | age<sup>2</sup> | 2 | age<sup>2</sup> | 2 |
| age × bio1<sub>t</sub> | 3 | age × PET<sub>t</sub> | 1 | age × bio5<sub>t</sub> | 4 |
| age × WAI<sub>p</sub> | 3 | age × prec.djf<sub>p</sub> | 1 | age × prec.jja<sub>p</sub> | 1 |
| bio1<sub>t</sub> × WAI<sub>p</sub> | 1 | PET<sub>t</sub> × prec.djf<sub>p</sub> | 1 | bio5<sub>t</sub> × prec.jja<sub>p</sub> | 1 |
| age × bio1<sub>t</sub> × WAI<sub>p</sub> | 3 | age × PET<sub>t</sub> ×<br>prec.djf<sub>p</sub> | 1 | age × bio5<sub>t</sub> ×<br>prec.jja<sub>p</sub> | 1 |

**Table S4.** Mean and standard deviation values for each phenotypic plasticity index. Analysis of the variance and post-hoc analyses adjusted by Tukey HSD were performed to test differences among developmental classes (DC).

|  |  | DC. 1 |  | DC. 2 |  | DC. 3 |  |
| --- | --- | --- | --- | --- | --- | --- | --- |
|  | P.I | mean | standard deviation | mean | standard deviation | mean | standard deviation |
| *P. nigra* | PP | 0.75a | 0.012 | 0.73b | 0.009 | 0.65c | 0.006 |
|  | CV | 0.48a | 0.015 | 0.45b | 0.011 | 0.36c | 0.006 |
| *P. pinaster* | PP | 0.86a | 0.011 | 0.85b | 0.001 | 0.84c | 0.007 |
|  | CV | 0.52a | 0.015 | 0.51b | 0.012 | 0.50c | 0.007 |
| *P. pinea* | PP | 0.98a | 0.000 | 0.98b | 0.000 | 0.95c | 0.000 |
|  | CV | 0.82a | 0.000 | 0.82b | 0.000 | 0.80c | 0.000 |

a, b and c (from the highest to the lowest) mean that mean values of phenotypic plasticity values are statistically different among developmental stages for each index analyzed.

**Table S5.**
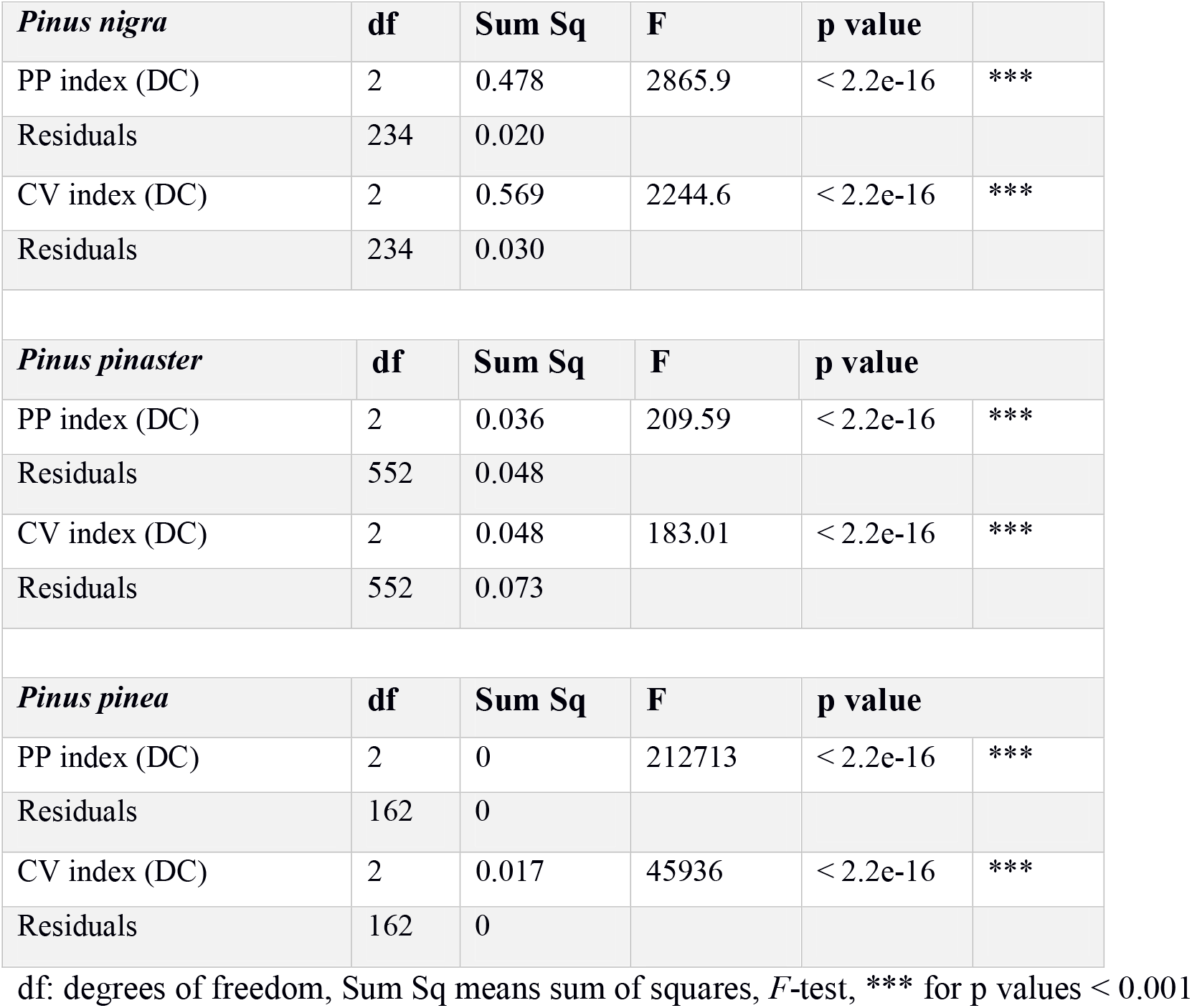
Results from the analysis of the variance to test phenotypic plasticity variation across developmental classes (DC) and for each pine species.

|  |  |  |  |  |  |
| --- | --- | --- | --- | --- | --- |
| *Pinus nigra* | df | Sum Sq | F | p value |  |
| PP index (DC) | 2 | 0.478 | 2865.9 | < 2.2e-16 | *** |
| Residuals | 234 | 0.020 |  |  |  |
| CV index (DC) | 2 | 0.569 | 2244.6 | < 2.2e-16 | *** |
| Residuals | 234 | 0.030 |  |  |  |
| *Pinus pinaster* | df | Sum Sq | F | p value |  |
| PP index (DC) | 2 | 0.036 | 209.59 | < 2.2e-16 | *** |
| Residuals | 552 | 0.048 |  |  |  |
| CV index (DC) | 2 | 0.048 | 183.01 | < 2.2e-16 | *** |
| Residuals | 552 | 0.073 |  |  |  |
| *Pinus pinea* | df | Sum Sq | F | p value |  |
| PP index (DC) | 2 | 0 | 212713 | < 2.2e-16 | *** |
| Residuals | 162 | 0 |  |  |  |
| CV index (DC) | 2 | 0.017 | 45936 | < 2.2e-16 | *** |

|  |  |  |
| --- | --- | --- |
| Residuals | 162 | 0 |

**Figure S1.**
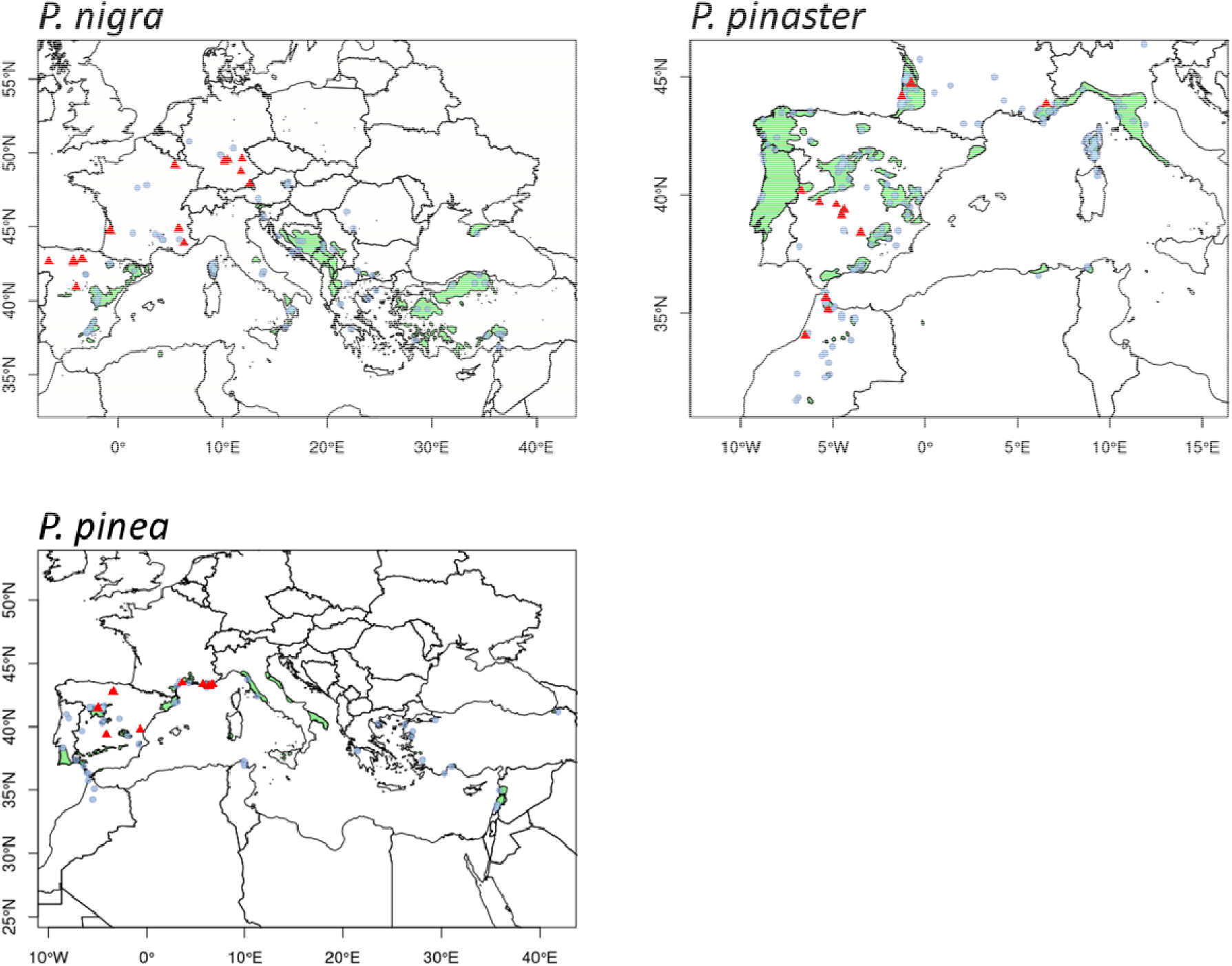
Maps showing the trials, provenances and natural distribution of the three species. Red triangles represent the common gardens (trials) and light blue circles the provenances. The light green area represents the natural distribution of the species according to EUFORGEN (http://www.euforgen.org/). Top left: *Pinus nigra*, top right: *Pinus pinaster*, bottom left: *Pinus pinea*. Adapted from Vizcaíno-Palomar *et al*., (2019).

**Figure S2.**
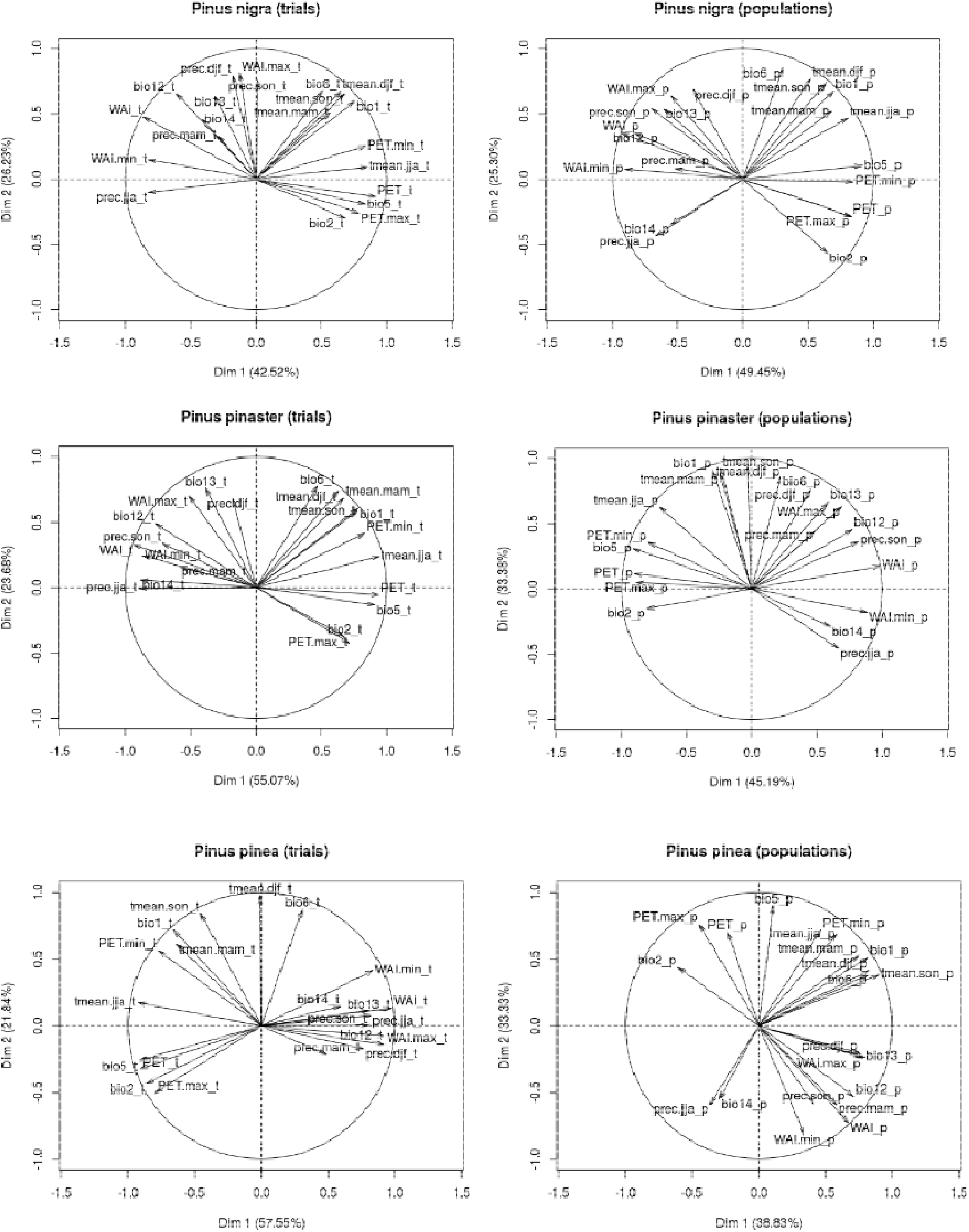
Plots of principal component analyses (PCA) for the short-term climate (clim_t_) and the long-term climate (clim_p_). These results are complementary to the linear mixed-effect model results shown in Appendix S2.

**Figure S3.**
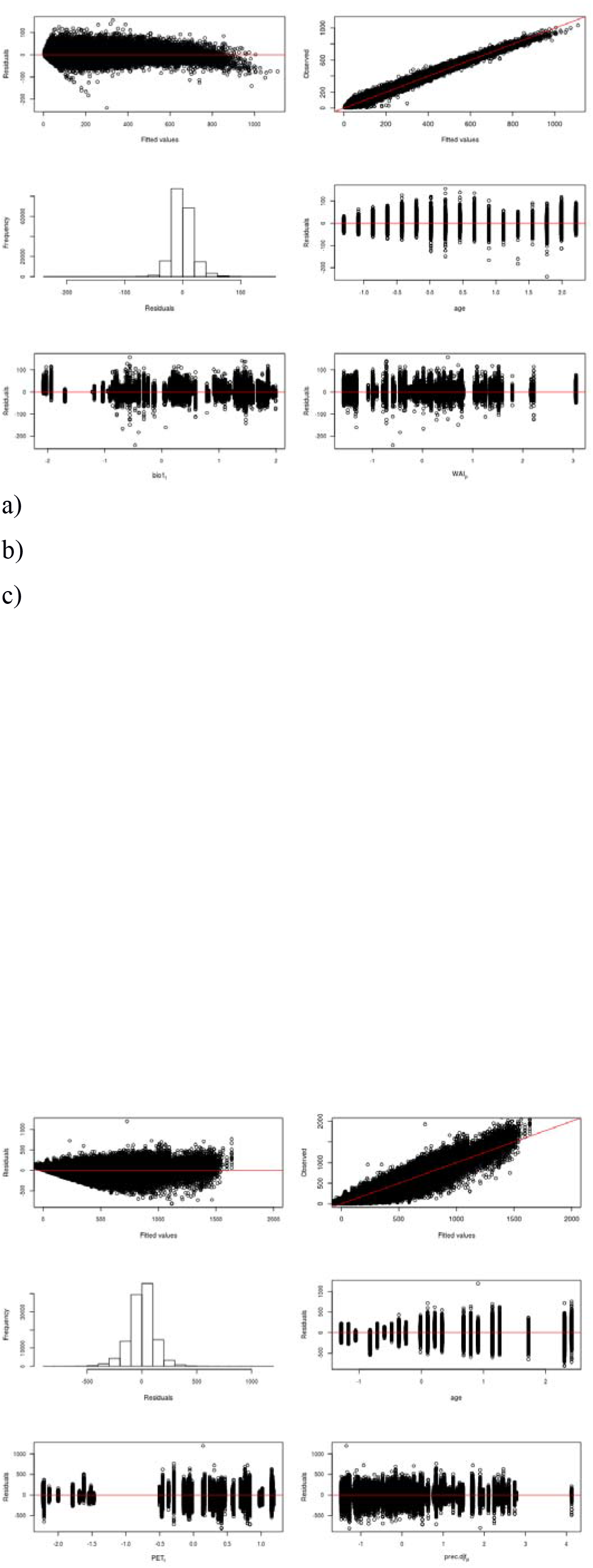
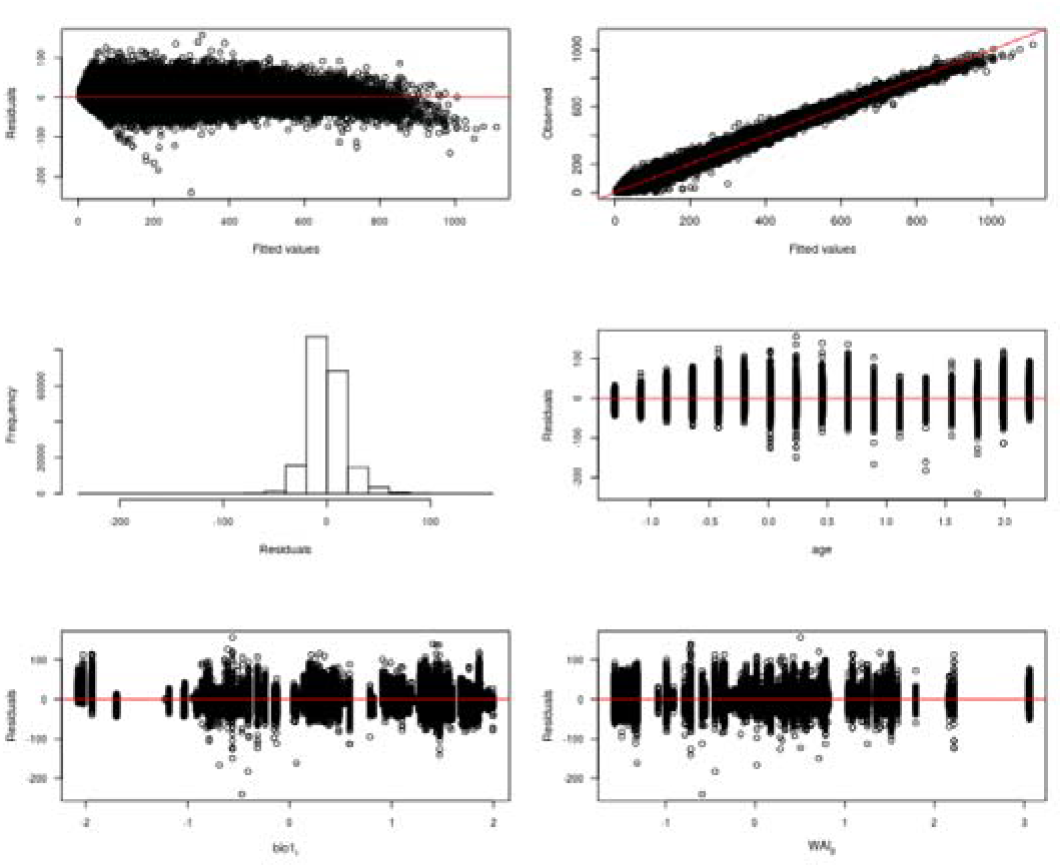
Plots of residuals of the best-supported model for tree height. Figures show the residuals across age, clim_t_ and clim_p_ in standardized values. a) *P. nigra,* b) *P. pinaster,* and c) *P. pinea*.

**Figure S4.**
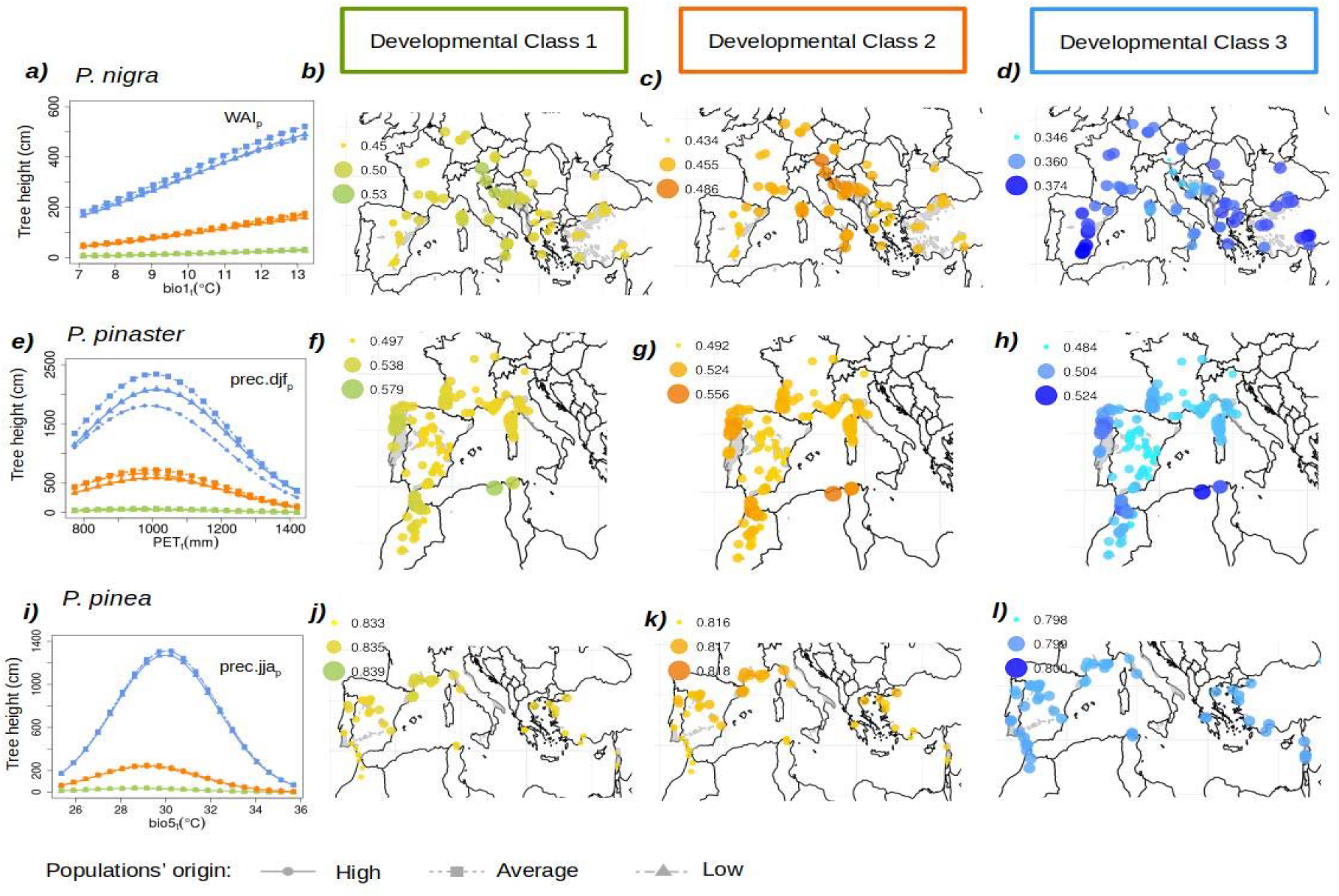
Populations’ phenotypic tree height responses across clim_t_ particularized for three populations’ origin (High, Average and Low in terms of clim_p_ values) and for the three developmental classes, DC, (Developmental Class 1: green, DC. 2: orange and DC. 3: blue) for a) *P. nigra*, e) *P. pinaster* and i) *P.pinea*. Values of the coefficient of variation index (CV) for the three developmental classes across the species natural distribution ranges are shown. DC. 1: b), f) and j), DC. 2: c), g) and k); and DC. 3: d), h) and i).

**Figure S5.**
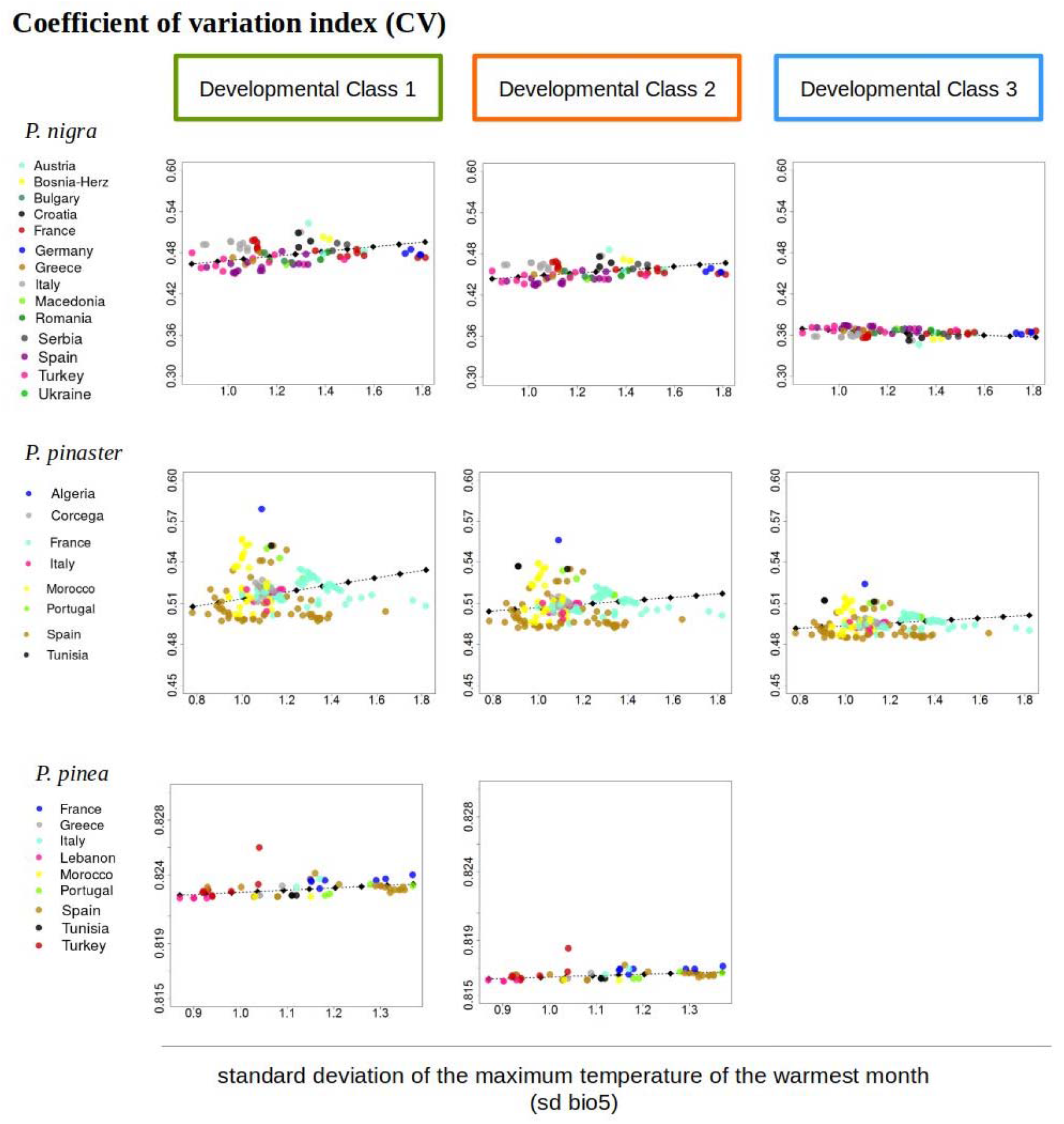
Phenotypic plasticity predictions of the CV index across the standard deviation of the maximum temperature of the warmest month (sd bio5) for the three developmental classes and pines species.

